# BIN1 regulates actin-membrane interactions during IRSp53-dependent filopodia formation

**DOI:** 10.1101/2022.03.21.485158

**Authors:** Laura Picas, Charlotte André-Arpin, Franck Comunale, Hugo Bousquet, Feng-Ching Tsai, Félix Rico, Paolo Maiuri, Julien Pernier, Stéphane Bodin, Anne-Sophie Nicot, Jocelyn Laporte, Patricia Bassereau, Bruno Goud, Cécile Gauthier-Rouvière, Stéphanie Miserey

**Affiliations:** Institut de Recherche en Infectiologie de Montpellier (IRIM), CNRS UMR 9004, University of Montpellier, Route de Mende, Montpellier, France; CRBM, University of Montpellier, CNRS, Montpellier, France; Institut Curie, PSL Research University, CNRS UMR 144, Paris, France; Institut Curie, PSL Research University, CNRS UMR 168, Paris, France; U1067 INSERM, Aix-Marseille Université, Marseille, France; Dipartimento di Medicina Molecolare e Biotecnologie Mediche, Università degli Studi di Napoli Federico II, Naples, Italy; Université Paris-Saclay, CEA, CNRS, Institute for Integrative Biology of the Cell (I2BC), Gif-sur-Yvette, France; Grenoble Alpes University, Inserm, U1216, Grenoble Institut Neurosciences, Grenoble, France; Department of Translational Medicine, IGBMC, U1258, UMR7104, Strasbourg University, Collège de France, Illkirch, France

**Author notes:** equal contribution (Cécile Gauthier-Rouvière, Stéphanie Miserey).

## Abstract

Amphiphysin 2 (BIN1) is a membrane and actin remodeling protein mutated in congenital and adult centronuclear myopathies. Here, we report an unexpected function of this N-BAR domain protein BIN1 in filopodia formation. We demonstrated that BIN1 expression is necessary and sufficient to induce filopodia formation. BIN1 is present at the base of forming filopodia and all along filopodia, where it colocalizes with F-actin. We identify that BIN1-mediated filopodia formation requires IRSp53, which allows its localization at negatively-curved membrane topologies. Our results show that BIN1 bundles actin in vitro. Finally, we identify that BIN1 regulates the membrane-to-cortex architecture and functions as a molecular platform to recruit actin-binding proteins, dynamin and ezrin, to promote filopodia formation.

## Introduction

Cellular morphologies powered by the actin cytoskeleton are essential to support developmental processes and cellular functions. Among the actin-driven structures produced by cells, filopodia assist relevant processes such as axonal guidance, establishing neuronal synapses, zippering of epithelial sheets, or myoblast fusion (Chen, 2011; Abmayr and Pavlath, 2012; Gallop, 2020). Filopodia are specialized 60-200 nm diameter finger-like structures made of paired actin filaments of a few to hundred microns in length (Mattila and Lappalainen, 2008; Svitkina, 2018). The remodeling of the membrane/actin interface plays a central role in the formation and shape maintenance of filopodia. This function is driven by modular proteins that connect actin microfilaments with the plasma membrane, typically via phosphatidylinositol 4,5-bisphosphate (PI(4,5)P_2_)-rich interfaces, as in the case of ezrin-radixin-moesin (ERM) domain proteins (Fievet et al., 2004; Garbett et al., 2013; Tsai et al., 2018), or the inverse-Bin1-Amphiphysin-Rvs (I-BAR) domain protein IRSp53 (Disanza et al., 2013; Prévost et al., 2015).

Bin/Amphiphysin/Rvs (BAR) domain proteins are multi-functional effectors characterized by a modular architecture consisting of an intrinsically curved membrane-binding BAR domain followed by auxiliary domains mediating protein-protein interactions and Rho GTPase signaling domains (Itoh and Decamilli, 2006; Suetsugu et al., 2010). One of the most prevalent auxiliary domains is the Src homology 3 (SH3) (Carman and Dominguez, 2018), and in many BAR domain proteins, it binds directly with cytoskeletal assembly factors and the dynamin GTPase (Itoh et al., 2005; Ferguson et al., 2009; Falcone et al., 2014; D’Alessandro et al., 2015). Despite the role of dynamin in endocytosis (Daumke et al., 2014), recent works support its central function as a multi-filament actin-bundling protein that propels cellular protrusions during myoblast fusion (Chuang et al., 2019; Zhang et al., 2020). Indeed, dense F-actin structures at the myoblast fusion site facilitate the formation of protrusions, allowing cell membrane juxtaposition that powers the mechanical forces required for cell-cell fusion and undergo myoblast differentiation (Sens et al., 2010; Kim et al., 2015). However, what precisely controls dynamin recruitment to initiate the formation of actin-rich protrusions at the plasma membrane remain unclear.

In skeletal muscle cells, the N-BAR domain protein BIN1/Amphiphysin 2 is a major binding partner of dynamin (Kojima et al., 2004; Nicot et al., 2007), and its expression appears highly induced during skeletal muscle differentiation (Wechsler-Reya et al., 1998; Lee et al., 2002). Indeed, the perturbed BIN1-dynamin interaction is the cause of centronuclear myopathies (CNMs), a heterogeneous group of inherited muscular disorders characterized by fiber atrophy and muscle weakness (Nicot et al., 2007; Fugier et al., 2011). Interestingly, the muscle-specific BIN1 isoform (BIN1 isoform 8), in contrast to the other BIN1/amphiphysin isoforms, displays a phosphoinositide (PI)-binding motif responsible for its targeting to the plasma membrane, mainly by interacting with PI(4,5)P_2_ (Lee et al., 2002). We previously showed that BIN1 locally increases the PI(4,5)P_2_ density to recruit dynamin on membranes selectively (Picas et al., 2014). Moreover, the PI domain controls a conformational switch that facilitates BIN1 SH3 domain interaction with dynamin (Kojima et al., 2004; Wu and Baumgart, 2014), and to other BIN1 partners such as the neuronal Wiskott-Aldrich syndrome (N-WASP) protein that regulates actin polymerization through the Arp2/3 complex (Yamada et al., 2009; Falcone et al., 2014; D’Alessandro et al., 2015). Furthermore, *in vitro* and *in cellulo* approaches have shown that neuronal and muscle-specific BIN1 isoforms directly bind F-actin (D’Alessandro et al., 2015; Dräger et al., 2017). However, it is unknown how BIN1 remodels the plasma membrane/actin interface and whether it fulfills a functional role in forming actin-rich protrusion.

Here, we report that BIN1 is a multi-functional platform to promote filopodia formation in myoblasts. Our results show that BIN1 assembles actin bundles *in vitro*. We also identified new BIN1-binding partners, including IRSp53 and ezrin, during filopodia formation. We show that BIN1 regulates the membrane-to-cortex mechanics, and it is required to recruit active phosphorylated ezrin at filopodia. Thus, we propose that BIN1 constitutes a functional membrane and actin-remodeling scaffold that, unexpectedly, as an N-BAR domain protein, orchestrates antagonistic membrane topologies at the cell cortex, such as the formation of filopodia.

## Results

### BIN1 expression is necessary and sufficient for filopodia-like structure formation

The association of different BIN1/Amphiphysin2 isoforms with F-actin and actin regulators such as the neuronal Wiskott-Aldrich syndrome (N-WASP) (Falcone et al., 2014; D’Alessandro et al., 2015; Dräger et al., 2017), prompt us to investigate if BIN1 proteins participate in actin-rich structure formation. Therefore, we expressed in HeLa cells GFP and several GFP-tagged amphiphysin isoforms: amphiphysin1 (typically associated with clathrin-coated structures) (McMahon and Boucrot, 2011), the largest BIN1 isoform (BIN1 iso1, expressed in neurons), and the muscle-specific isoform BIN1 (BIN1 iso8), which contains the in-frame exon 11 encoding a polybasic motif binding phosphoinositides (PIs) (Prokic et al., 2014), and stained F-actin. We observed that BIN1 iso8 drastically enhanced filopodia-like structure density, whereas BIN1 iso1 and Amphiphysin1 led to a low to null increase in filopodia-like structure density, respectively (Fig. 1A).

**Figure 1.**
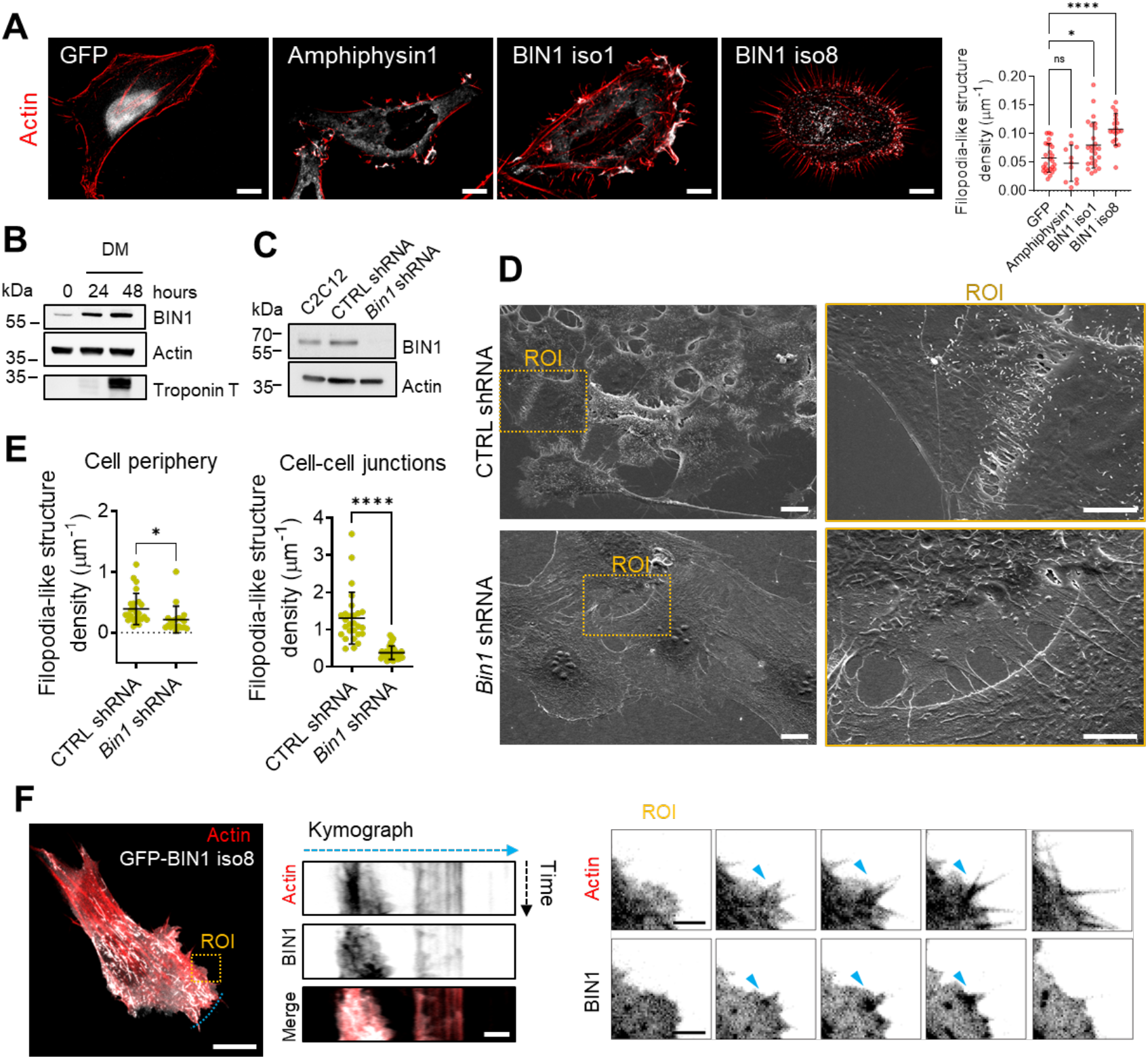
BIN1 expression promotes the formation of filopodia-like structures. **A)** Wild-field deconvoluted images of HeLa cells transfected with GFP or GFP-tagged Amphiphysin1, BIN1 iso1 and BIN1 iso8 (in gray) and stained for F-actin (phalloidin, red). Scale bar, 10 µm. Quantification of the filopodia-like structure density of HeLa cells expressing GFP or GFP-tagged Amphiphysin1, BIN1 iso1 and BIN1 iso8. Number of cells: n =26, 11, 25 and 18, respectively from three independent experiments. **B)** Western-blot analysis of the endogenous expression of actin, BIN1, and Troponin T in C2C12 myoblasts in growth medium (0 hours, undifferentiated) and grown in differentiation medium (DM) at 24h and 48h. **C)** Westernblot analysis of the endogenous expression of actin and BIN1 on parental, CTRL shRNA (i.e. Luciferase) and *Bin1* shRNA C2C12 myoblast cells. **D**) Scanning electron microscopy images of CTRL shRNA and *Bin1* shRNA C2C12 myoblasts cultured in growth factor-containing medium. Scale bar, 10 µm and 5 µm for the magnified images. **E)** Quantification of the number of filopodia-like structures per cell at the cell periphery and at intercellular junctions measured from the SEM images. Cells: *n*=25, 19 and 27, 38 for CTRL and *Bin1* shRNA cells at the cell periphery and cell-cell junctions, respectively. **F**) Snap-shots of spinning disk life cell imaging (500 msec exposure, during 60s. Total time acquisition = 120s) of C2C12 myoblasts co-transfected with Lifeact-mCherry (red) and GFP-BIN1 iso8 (gray) at t = 0s. Kymograph analysis along the blue dashed line in the corresponding image at t = 0s, highlight the recruitment and binding of BIN1 iso8 to filopodia-like structures. Scale bar, 10 µm. Scale bar in kymograph and ROI, 2 µm. Bottom, representative time-lapse snapshots from the ROI region showing the localization of BIN1 iso8 on F-actin during filopodia-like structure formation in C2C12 cells, as highlighted by the blue arrowheads. Error bars represent s.d.; ANOVA test: n.s > 0.1, * P < 0.1, **** P < 0.0001.

To inquiry into the molecular mechanisms of BIN1-mediated filopodia-like structure formation, we turned out to myoblast cells because actin protrusions have been implicated in the adhesion and fusion of muscle cells (Segal et al., 2016; Zhang et al., 2020) and BIN1 expression increases during myoblast differentiation (Fig. 1B) (Lee et al., 2002). To this end, we generated C2C12 myoblast expressing specific short interfering RNA (shRNA) against *Bin1* or luciferase (CTRL shRNA) to ensure a homogenous knock-down of *Bin1* in the myoblast cell population. In undifferentiated *Bin1* shRNA C2C12 cells, depletion of BIN1 expression was almost complete (Fig. 1C). We then performed a detailed analysis of the cellular morphology of control and C2C12 *Bin1* shRNA myoblasts by scanning electron microscopy (SEM) either under proliferative conditions and at 80-90% confluence to favor the formation of cell-cell junctions preceding myoblast differentiation and fusion. Furthermore, BIN1 depletion was associated with reduced membrane extensions between cells and a smoother dorsal plasma membrane (Fig. 1D-contacting cells and Fig. S1A-isolated cells). This phenotype was evident at the cell periphery and cell-cell junctions, indicating that BIN1 might participate in adjoining myoblasts’ intercellular zippering. Conversely, we observed a dense interface of filopodia-like membrane protrusions interconnecting two or more adjacent myoblasts in control cells. In *Bin1* knock-down cells, the protrusion densities are significantly decreased (3.6-fold reduction in filopodia-like structure density, Fig. 1E). These results show that in myoblasts, BIN1 depletion affects the morphology and the number of filopodia-like structure at cell-cell junctions and at the cell periphery (i.e., away from intercellular junctions).

Next, we analyzed BIN1 localization by multi-color life-cell imaging of C2C12 cells co-expressing Lifeact-mCherry and BIN iso8. We showed that at the cell periphery BIN1 co-localized with actin at filopodia-like structures (Fig. 1F). Furthermore, kymograph analysis of the GFP-BIN1 signal showed that it often precedes that of Lifeact-mCherry (blue arrowheads, Fig. 1F). Moreover, BIN1 signal is present during the initiation, extension, and retraction of filopodial actin filaments (Fig. 1F).

Altogether, these data show that BIN1 expression promotes filopodia-like structure formation and that BIN1 knockdown decreases these F-actin rich structures.

### BIN1 promotes actin bundling

BIN1 iso1 was previously reported to bundle actin *in vitro* (D’Alessandro et al., 2015; Dräger et al., 2017) and its N-BAR and SH3 domains share a strong homology with BIN1 iso8 (Prokic et al., 2014). Thus, we sought to determine if the muscle-specific BIN1 iso8 also has the ability to associate and remodel F-actin. First, we performed a high-resolution analysis of BIN1 and F-actin using structured illumination microscopy (SIM) imaging in C2C12 cells. The endogenous BIN1 staining (Fig. 2A) or expression of GFP-BIN1 iso8 (Fig. 2B) revealed that BIN1 is localized at the base of filopodia structures and displays a discontinuous labeling along these F-actin-rich structures. Consequently, we determined whether BIN1 localization in filopodia-like structures is related to its association with actin. Thus, we performed co-immunoprecipitation experiments with GFP-tagged amphiphysin1, BIN1 iso1, and BIN1 iso8 (Fig. 2C). Immuno-blotting showed that *in cellulo,* only BIN1 iso8 appears associated with actin. Next, we determined the contribution of the BIN1 N-terminal N-BAR and the C-terminal SH3 domain for its association with actin *in cellulo* (Fig. 2D). To this end, we tested by co-immunoprecipitation the interaction with actin of the BIN1 D151N mutant, which carries a mutation in the N-BAR domain, which interferes with the curvature-sensing abilities and capacity to oligomerize (Wu et al., 2014), and of the BIN1 ΔSH3 mutant, which displays a truncated SH3 domain shown to prevent its interaction with cellular effectors (Nicot et al., 2007) or N-WASP (Falcone et al., 2014). As shown in Fig. 2D, both mutants associate with actin *in cellulo*.

**Figure 2.**
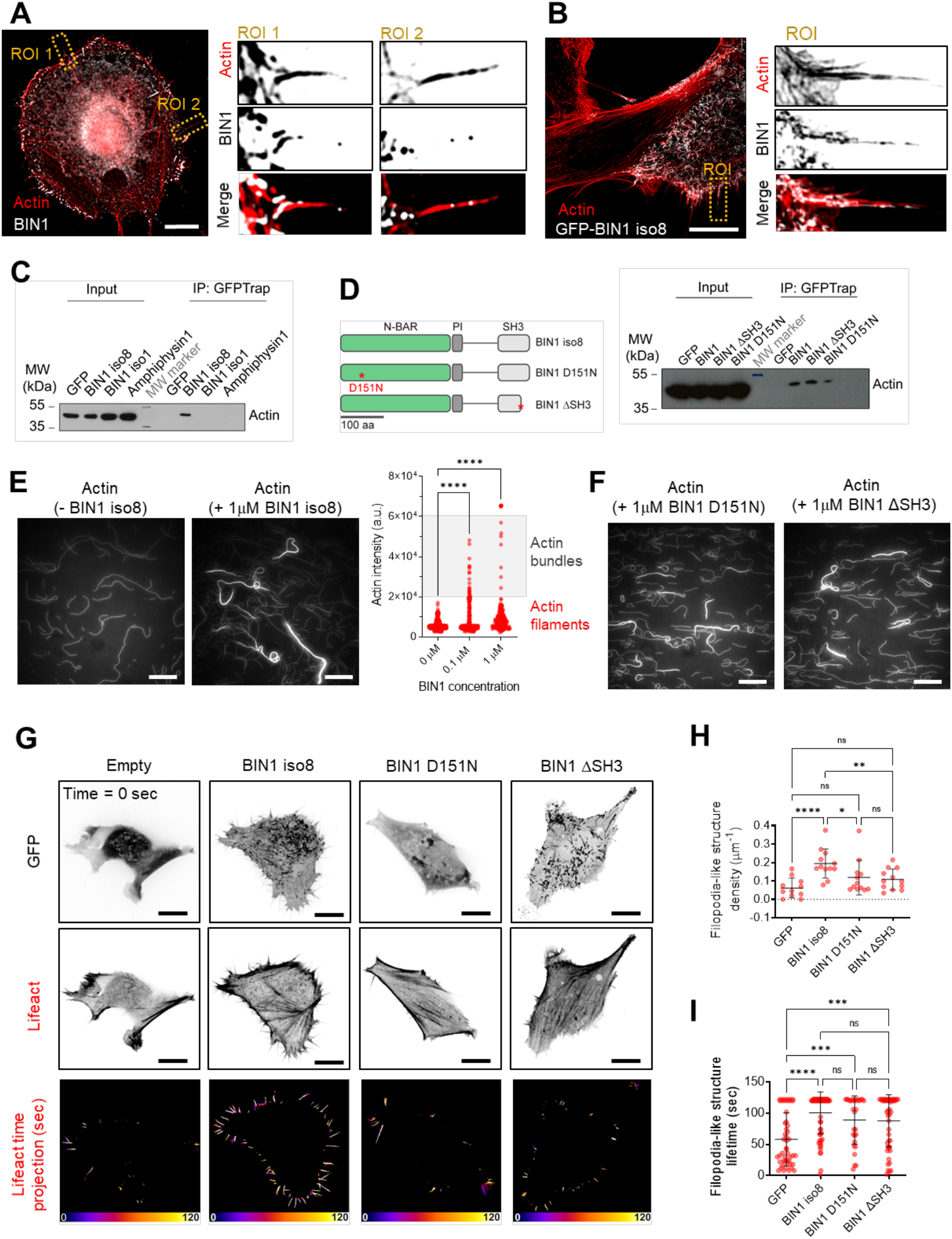
BIN1 interacts with F-actin *in vitro* and *in cellulo*. **A)** C2C12 myoblasts stained for endogenous BIN1 (C99D antibody, gray) and F-actin (phalloidin, red). Magnified images of two representative regions of interest (ROIs). Scale bar, 10 µm. **B)** SIM images of C2C12 myoblasts expressing GFP-BIN1 (gray) stained for F-actin (phalloidin, red) and magnified images of two representative ROI in the corresponding image. Scale bar, 10 µm. **C)** GFP pull-downs using extracts from HeLa cells expressing either GFP, or GFP-BIN1 iso8, GFP-BIN1 iso1 and GFP-Amphiphysin1. Actin was detected by western blotting. HeLa cells were used to provide an unbiased cellular context. IP = immunoprecipitate. **D)** Domain representation (N-BAR, phosphoinositide-binding motif, PI, and SH3 domains) of BIN1 iso8 full-length. Stars highlight the D151N mutant at the N-BAR and the stop codon of the BIN1 ΔSH3 mutant. GFP-Trap pull-downs using extracts from C2C12 myoblasts expressing either GFP alone or fused with BIN1 iso8, BIN1 ΔSH3 or BIN1 D151N. Actin was revealed by western blotting. **E)** *In vitro* assay with purified actin and BIN1 iso8. TIRF images showing pre-polymerized F-actin filaments in the absence (- BIN1) or in the presence of BIN1 (+ 1 µM BIN1 iso8) and the corresponding quantification of the BIN1 bundling activity as denoted by the actin intensity in each condition. Scale bar, 20 µm. **F)** *In vitro* assay with purified actin and BIN1 mutants BIN1 ΔSH3 or BIN1 D151N. TIRF images showing pre-polymerized F-actin filaments in the presence of 1 µM BIN1 mutants. Scale bar, 20 µm. **G)** Snap-shots of spinning disk life cell imaging at t = 0 of C2C12 cells cotransfected with mCherry-Lifeact and either GFP (empty), GFP-BIN1 iso8, GFP-BIN1 D151N or GFP-BIN1 ΔSH3 (inverted LUT images of the actin and GFP signal). Time projection of the filopodia-like structure tracks (500 msec exposure, during 60s. Total time acquisition = 120s) obtained from of the corresponding Lifeact images. Scale bar, 10 µm. Fire LUT color scale is 120s. **H**) Quantification of F-actin-rich filopodia-like structure density; Cells: *n*=11, 12, 12 and 12 for GFP, BIN1 iso8, D151N and ΔSH3 mutants, respectively. **I)** Filopodia lifetime; *n*=253, 1743, 711 and 785 for GFP, BIN1 iso8, D151N and ΔSH3 mutants, respectively. Error bars represent s.d.; ANOVA test: n.s > 0.1, * P < 0.1, ** P < 0.01, *** P < 0.001, **** P < 0.0001.

The co-localization and association of BIN1 iso8 with actin *in cellulo* prompted us to determine the potential ability of BIN1 iso8 to remodel F-actin *in vitro* using an assay based on purified actin and BIN1. We showed that BIN1 promotes actin filamentbundling (Fig. 2E). Remarkably, the number of actin bundles increased with increasing BIN1 concentrations, although we already observed a bundling effect at 0.1 μM of BIN1 with 1 μM of actin, as compared to the actin-bundling activity reported for the BIN1 iso1 (≥ 0.25 μM according to (Dräger et al., 2017)). Furthermore, we detected the similar actin-bundling activity by adding each of the BIN mutants separately (Fig. 2F). Altogether, these data show that mutations at the N-BAR and SH3 domain truncation do not perturb the F-actin association and bundling ability of BIN1.

Next, we analyzed the effect of the N-BAR and SH3 domain of BIN1 on filopodia-like structure formation *in cellulo* (Fig. 2G-I Fig. S1B). Therefore, GFP, GFP-BIN1 iso8, or GFP-tagged BIN1 D151N and BIN1 ΔSH3 mutants were co-expressed with Lifeact-mCherry to monitor the density and lifetime of actin-rich protrusions. As expected, expression of GFP-BIN1 iso8 in C2C12 cells increased filopodia density compared to control conditions (GFP alone). However, we did not detect a significant increase in their density with the D151N or ΔSH3 BIN1 mutants (Fig. 2G and H). Nevertheless, no significant differences were observed in the filopodia lifetime (i.e., the time until filopodia collapse) between BIN1 wild-type expression (∼ 2-fold increase in the filopodia lifetime) and the two BIN1 mutants (∼ 1.5-fold increase) (Fig. 2I), in agreement with their similar association to and bundling of F-actin (Fig. 2D-F). These data indicate that the N-BAR and SH3 domains are likely to contribute to BIN1-induced filopodia-like structure formation, possibly favoring the association with BIN1-binding partners.

### IRSp53 is required for BIN1-mediated filopodia formation

To evaluate the contribution of potential BIN1 partners in the formation of F-actin-rich structures, we performed a proteomic analysis using GFP-BIN1 iso8 to identify its binding partners (Fig. S2A). We found already known BIN1 partners (in grey Fig. S2A): dynamin 1 and 2 (Lee et al., 2002; Nicot et al., 2007) and actin (this study and D’Alessandro et al., 2015), and unknown partners (in yellow Fig. S2A) such as IRSp53 and ezrin. Interestingly, IRSp53 is involved in the Cdc42-dependent formation of filopodia (Scita et al., 2008; Lim et al., 2008; Disanza et al., 2013). Therefore, we confirmed by immunofluorescence that endogenous IRSp53 and BIN iso8 colocalize at actin-rich filopodia-like structures on C2C12 myoblasts (Fig. 3D). Next, we determined the role of IRSp53 in BIN1-mediated filopodia formation in C2C12 cells (Fig. 3E-G). As expected, IRSp53 knock-down in C2C12 cells led to a decrease in filopodia-like structure density. While the expression of GFP-BIN1 iso8 in CTRL siRNA C2C12 promoted an increase in filopodia formation, this was not the case under IRSp53 knock-down (Fig. 3G), therefore, suggesting that BIN1-mediated filopodia-like structure formation requires the inverted-BAR (I-BAR) protein IRSp53.

**Figure 3.**
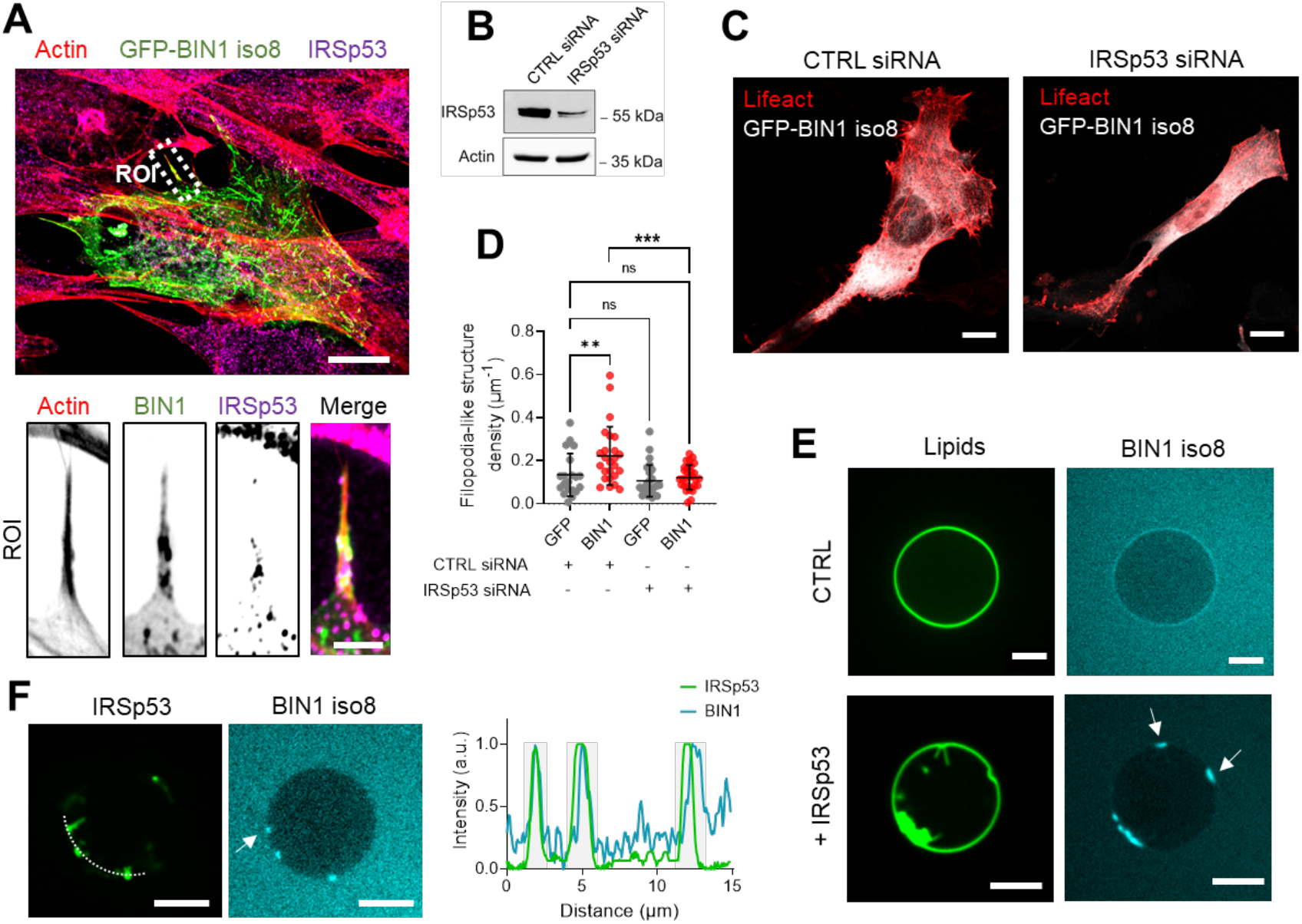
BIN1 associates with IRSp53 on membrane protrusions. **A)** Confocal/airyscan images of C2C12 cells transfected with GFP-BIN1 iso8 (in green) and stained for endogenous IRSp53 (cyan) and F-actin (phalloidin, red). Magnified image of the corresponding ROI. Scale bar, 10 µm. Scale bar ROI, 2 µm. **B)** Western-blot analysis of the endogenous expression of actin and IRSp53 on CTRL siRNA (i.e. Luciferase) and IRSp53 siRNA C2C12 cells. **C)** Z-projected confocal images showing the actin organization of CTRL siRNA and IRSp53 siRNA C2C12 cells co-transfected with GFP-BIN1 iso8 (gray) and Lifeact-mCherry (red). **D)** Quantification of filopodia-like structure density; *n*=22, 26, 26 and 27 for CTRL siRNA and IRSp53 siRNA C2C12 cells co-transfected with either GFP or GFP-BIN1 and Lifeact-mCherry respectively. Error bars represent s.d.; ANOVA test: n.s > 0.1, ** P < 0.01, *** P < 0.001. **E)** Representative confocal images of 0.1 μM BIN1 binding (cyan) to control (only BIN1) or in the presence of 0.3 μM IRSp53-induced tubules on PI(4,5)P2-containing GUVs (green). Scale bar, 5 µm. **F)** Representative confocal images of 0.1 μM BIN1 binding (cyan) to PI(4,5)P2-containing GUVs (dark) in the presence of 0.3 μM IRSp53 (green). Profile analysis of BIN1 and IRSp53 signal along the dashed line in the corresponding image. Scale bar, 5 µm.

The topology at the neck of filopodia is compatible with the binding of proteins with positive curvature BAR-domain sensors like BIN1 (Suetsugu et al., 2010). Therefore, we inquire whether BIN1 iso8 can be recruited to negatively-curved membranes induced by IRSp53 using an *in vitro* reconstituted assay and confocal microscopy (Fig. 3E), as previously reported (Inamdar et al., 2021). We generated 5% mole PI(4,5)P_2_-containing giant unilamellar vesicles (GUVs) doped with Oregon Green 488 DHPE, as PI(4,5)P_2_ is a central phosphoinositide for BIN1 and IRSp53 membrane interactions (Mattila et al., 2007; Picas et al., 2014). As expected, the addition of the fulllength IRSp53 led to the formation of inward membrane tubes on PI(4,5)P_2_-GUVs. The addition of recombinant alexa fluor A647-labeled BIN1 iso8 showed that it accumulates to membrane indentations that are generated by IRSp53 (Fig. 3E). To confirm that the presence of IRSp53 mediates BIN1 localization to the base of inward membrane tubes, we generated dark PI(4,5)P_2_-containing GUVs and monitored the co-localization of recombinant alexa fluor A488-labeled IRSp53 and alexa fluor A647-labeled BIN1 iso8 (Fig. 3F). Our results confirmed that IRSp53 allows BIN1 recruitment at the base of negatively-curved membrane structures.

### BIN1-mediated recruitment of ezrin participates in filopodia formation

Interestingly, our proteomic study identified novel BIN1 interactors, such as the members of the ezrin-radixin-moesin (ERM) protein family (Fig. S2A), previously found to accumulate at the shaft of filopodia and to co-localize with IRSp53 on microvilli (Gandy et al., 2013; Garbett et al., 2013). To study if specificity exists between different BIN isoforms and the two ERM members ezrin and moesin, we performed co-immunoprecipitation experiments using several amphiphysin isoforms: amphiphysin1, BIN1 iso1 and BIN1 iso8. Our results showed that only BIN1 iso8 associated with ezrin *in cellulo* (Fig. 4A), while both BIN1 iso1 and iso8 isoforms associate with moesin (Fig. S2B). We also found that only the BIN1 D151N mutant, but not the ΔSH3 mutant, associates with ezrin (Fig. 4B), suggesting that the SH3-containing C-terminal domain of BIN1 mediates the interaction with ezrin.

**Figure 4.**
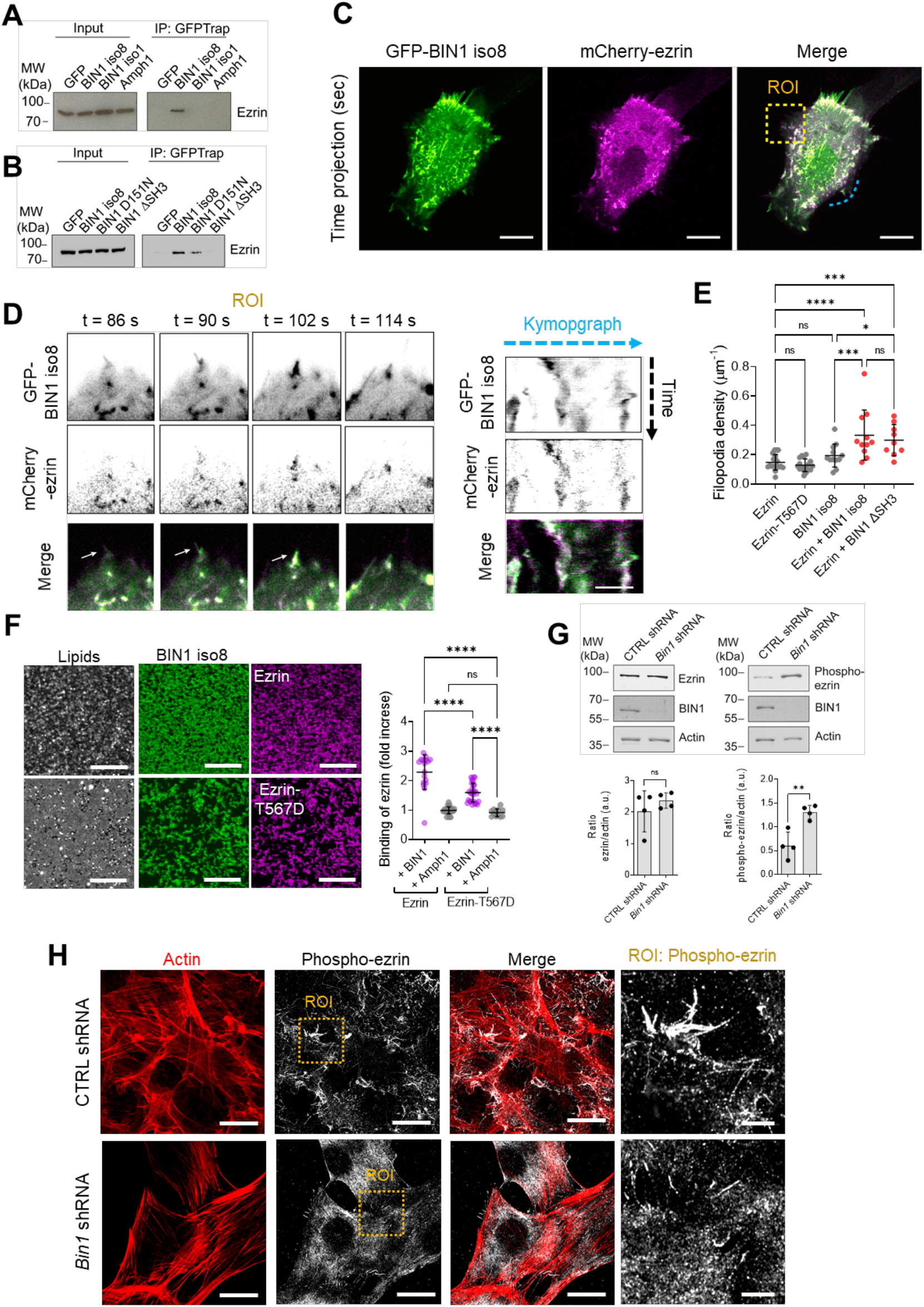
BIN1 regulates ezrin localization on membranes. **A)** GFP-Trap pull-downs from extracts of HeLa cells transfected with plasmids encoding GFP, GFP-tagged amphiphysin1, BIN1 iso1 and BIN1 iso8. Ezrin was revealed by western-blotting. **B)** GFP-Trap pull-downs using extracts from C2C12 myoblasts expressing either GFP alone or fused with BIN1 iso8, BIN1 ΔSH3 or BIN1 D151N. Ezrin was revealed by western-blotting. **C)** Time projected spinning disk movies (500 msec exposure, during 60s. Total time acquisition = 120s) of C2C12 cells co-expressing mCherry-ezrin (magenta) and GFP-BIN1 iso8 (green). **D)** Time-lapse snapshots from the corresponding yellow ROI showing the co-localization of BIN1 and ezrin during filopodia-like structure formation (white arrows). Kymograph analysis along the blue dashed line. Scale bar, 10 µm. Scale bar in kymographs, 1 µm. **E**) Quantification of filopodia-like structure density; *n*=19, 17, 12, 10, and 10 for ezrin, ezrin-T567D, BIN1, ezrin + BIN1 iso8, and ezrin + BIN1 ΔSH3 mutant, respectively. n = number of cells from live cell imaging experiments. Error bars represent s.d.; ANOVA test: n.s > 0.1, ** P < 0.01, *** P < 0.001. **F)** Confocal images of the colocalization of recombinant BIN1-Alexa647 (magenta), ezrin or phospho-mimetic ezrin (T567D) tagged with Alexa488 (green) and TopFluor-TMR-PI(4,5)P2 (cyan) on supported lipid bilayers containing 5% of PI(4,5)P2. Scale bar, 3 µm. Fold increase in the binding of recombinant ezrin and ezrin-T567D on supported lipid bilayers containing 5% PI(4,5)P2 in the presence of BIN1 or amphiphysin1 (Amph1). **G)** Western-blot analysis of the endogenous expression of ezrin and phosphorylated ezrin (phospho-ezrin), BIN1 and actin on cell extracts from CTRL (blue) and *Bin1* shRNA (yellow) C2C12 myoblasts, and the corresponding quantification of the signal of each protein normalized by actin. **H)** CTRL and *Bin1* shRNA C2C12 myoblasts cultured in growth factor-containing medium and stained for endogenous phosphorylated ezrin (phospho-ezrin, yellow) and F-actin (phalloidin, cyan). Inset, high-magnification images showing the localization of endogenous phosphorylated ezrin. Maximum intensity projected airyscan images. Scale bar, 10 µm and 5 µm, respectively.

Next, we tested the association of BIN1 and ezrin during filopodia formation. We performed multi-color time-lapse live-cell imaging of C2C12 cells co-expressing mCherry-ezrin together with either GFP-BIN1 iso8 and the ΔSH3 mutant that, according to our biochemical assay does not associate with ezrin (Fig. 4C and Fig. S2C). Spinningdisk images showed that BIN1 and ezrin co-localize at the cell periphery (Fig. 4C-D). Detailed analysis of the dynamics of ezrin and BIN1 also showed that both proteins co-localized during filopodia formation (Fig. 4D). In addition, co-expression of ezrin with BIN1 potentiated the formation of filopodia compared to BIN1 alone (Fig. 4E), suggesting that BIN1 and ezrin might cooperate in this process. Conversely, an absence of co-localization and of effect on filopodia density was observed between ezrin and the BIN1 ΔSH3 mutant, indicating that BIN1-ezrin mediated filopodia-like structure formation might require ezrin association to BIN1 via the SH3 domain of BIN1 (Fig. S2C).

Ezrin displays different conformational states, including a cytosolic closed-conformation, which results from the intramolecular interaction of the N-terminal FERM domain with the C-terminal ERM-associated domain (C-ERMAD), a membrane-bound opened-conformation, which requires a sequential activation through the interaction with PI(4,5)P_2_, a phosphorylated conformation, which displays a phosphorylation of a conserved threonine in the actin-binding site of the C-ERMAD (T567, in ezrin) and finally, the interaction of the C-ERMAD with F-actin (Fievet et al., 2004). Interestingly, we previously showed that BIN1 clusters PI(4,5)P_2_ to recruit PI(4,5)P_2_-interacting downstream partners on membranes (Picas et al., 2014). Therefore, we analyzed if PI(4,5)P_2_, a key phosphoinositide for the recruitment of actin regulatory proteins promoting actin polymerization and filopodia formation at the plasma membrane (Senju et al., 2017; Senju and Lappalainen, 2019) could participate in ezrin membrane recruitment by BIN1. We confirmed that filopodia positive for BIN1 are highly enriched in PI(4,5)P_2_ (Fig. S2D). Next, using an *in vitro* reconstituted assay consisting of supported lipid bilayers doped with 5% PI(4,5)P_2_, we quantified the binding of recombinant wild-type ezrin and its phospho-mimetic form (ezrin-T567D) in the presence of recombinant BIN1 iso8 (Fig. 4F). As a control we used amphiphysin 1, a protein that does not interact with ezrin, but that mediates PI(4,5)P_2_ clustering (Picas et al., 2014). We estimated the relative binding of ezrin, wild type or T567D, from the ratio between the intensity of ezrin proteins bound on membranes in the presence of BIN1 iso8 or of amphiphysin1, normalized by the intensity of ezrin proteins in the absence of BIN1 or amphiphysin1. We obtained a ∼ 2.5-fold and 1.5-fold increase in the relative membrane binding of ezrin and ezrin-T567D, respectively, in the presence of BIN1 and PI(4,5)P_2_ (Fig. 4F). In agreement with the results in Fig. 4A, amphiphysin1 did not affect ezrin binding to membranes. Altogether these observations suggest that BIN1-mediated ezrin recruitment on membranes is probably mediated by proteinprotein interactions (Fig. 4B) on PI(4,5)P_2_ interfaces.

To further understand the molecular mechanisms of BIN1/ezrin interaction at the cell cortex, we investigated the impact of *Bin1* knock-down on ezrin and phosphorylated ezrin (phospho-ezrin) expression (Fig.4G) and localization (Fig.4H). We observed a two-fold increase of phosphorylated ezrin in *Bin1* shRNA C2C12 cells, but no modification of the total pool of ezrin (Fig. 4G). In addition, we observed that phospho-ezrin appears enriched at filopodia in control C2C12 cells (Fig. 4H). However, such enrichment was not observed in C2C12 *Bin1* shRNA myoblasts. These results indicate that although the total level of phospho-ezrin is increased, the cellular organization of phospho-ezrin appears affected in *Bin1* knock-down myoblasts. This suggests that BIN1 might regulate the localization and activation of ezrin at the plasma membrane.

### BIN1 regulates the cell cortex architecture of myoblast

In myoblast cells, BIN1 expression increases during differentiation (Fig. 5A) (Lee et al., 2002; Nicot et al., 2007), and its inhibition is associated with impaired myotube formation (Wechsler-Reya et al., 1998; Lee et al., 2002), data that we also confirmed (Fig. S3). Furthermore, we observed that the expression of different differentiation markers was not affected in *Bin1* shRNA C2C12 (Fig. S3), suggesting a potential role of BIN1 during the fusion stages of myoblast cells. Previous works showed that the SH3 domain of BIN1 mediates the interaction with its downstream partner dynamin (Kojima et al., 2004; Royer et al., 2013), a GTPase well known to bundle actin filaments and having an important role during filopodia formation promoting myoblast fusion *in vivo* (Gu et al., 2010; Yamada et al., 2016; Chuang et al., 2019; Zhang et al., 2020). Therefore, we confirmed by immunofluorescence that endogenous BIN1 colocalizes with GFP-dynamin2 at actin-rich structures on C2C12 cells (Fig. 5A). In contrast, the density and morphology of dynamin2-positive actin-rich protrusions appeared affected in C2C12 *Bin1* shRNA myoblasts (Fig. 5B-C). Collectively, supporting that, in addition to their shared role in membrane remodeling, BIN1 and its partner dynamin2 can also interact on actin-rich protrusions.

**Figure 5.**
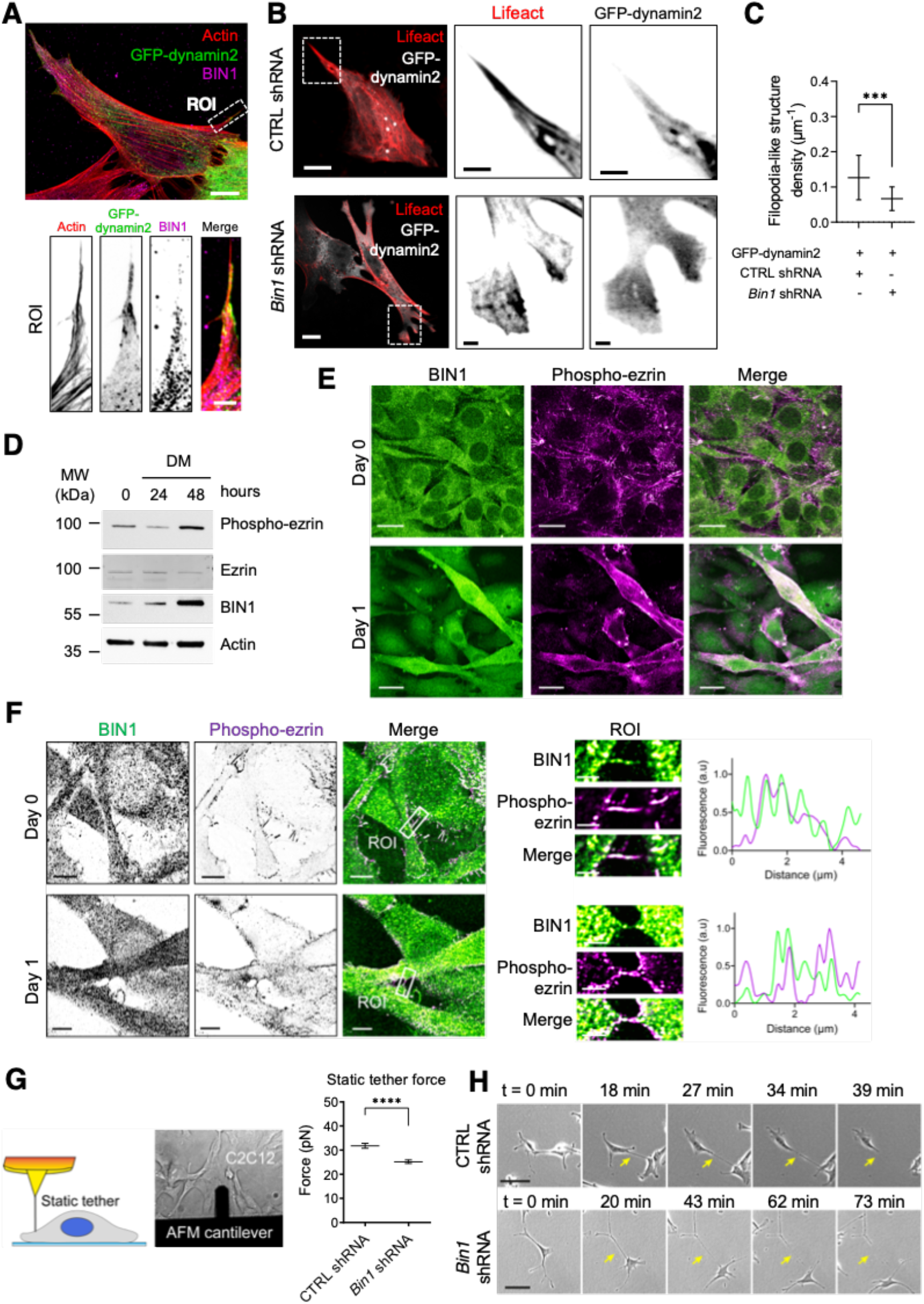
BIN1 regulates the cell cortex architecture. **A)** Confocal/airyscan images of C2C12 cells transfected with GFP-dynamin2 (in green) and stained for endogenous BIN1 (magenta) and F-actin (phalloidin, red). Magnified image of the corresponding ROI. Scale bar, 10 µm. Scale bar ROI, 2 µm. **B)** Z-projected confocal images showing the actin organization of CTRL shRNA and *Bin1* shRNA C2C12 cells co-transfected with GFP-dynamin2 (gray) and Lifeact-mCherry (red). **C)** Quantification of filopodia-like structure density; number of cells, *n*=20 and 17 for CTRL shRNA and *Bin1* shRNA C2C12 cells co-transfected with GFP-dynamin2 and Lifeact-mCherry respectively. Error bars represent s.d.; t-test: *** P < 0.001. **D)** Western-blot analysis of the endogenous expression of BIN1, ezrin and phosphorylated ezrin (phospho-ezrin) in C2C12 myoblasts under proliferative conditions (Day 0) or at 24h and 48h in differentiation medium (Day 1 and Day 2, respectively). **E)** Maximum intensity projected confocal images of proliferative myoblasts (Day 0) or at 24h in differentiation medium (Day 1) stained for endogenous BIN1 (C99D antibody, green) and phosphorylated ezrin (phospho-ezrin, magenta). Scale bar, 20 µm. **F)** Airyscan images of C2C12 myoblasts at day 0 and day 1 stained for endogenous BIN1 (green) and phospho-ezrin (magenta). Cross-section along a representative filopodia-like structure (BIN1 in green, phospho-ezrin in magenta) highlighted by the white box and magnified in the ROI image. Scale bar is 5 µm, and 2 µm for the inset. **G)** Schematic representation of a plasma membrane tether pulling assay using an AFM cantilever coated with poly-L-lysine on adherent C2C12 cells. Representative bright field image of the corresponding experimental setup. Static tether force (pN) obtained with CTRL (blue) and *Bin1* shRNA (yellow) C2C12 myoblasts. t-test: P < 0.0001. **H)** Snapshots of bright field images of CTRL and *Bin1* shRNA C2C12 myoblasts cultured in growth factor-containing medium at different time points. Yellow arrow highlights the retraction of retraction fibers formed during cell migration. Scale bar, 50 µm.

Interestingly, ezrin, one of our BIN1-interacting partners during filopodia formation, was recently shown to be involved in myoblast differentiation and fusion (Zhang et al., 2023). Therefore, to investigate if the association of BIN1/ezrin on filopodia is maintained during myoblast differentiation, we analyzed the expression (Fig. 5D-E) and co-localization of endogenous BIN1 and the active phosphorylated form of ezrin (phospho-ezrin) in proliferating C2C12 cells (Day 0), i.e. ≤ 80-90% cell confluency in growth medium, and at 1 day of differentiation (Day 1), i.e. 24h in differentiation medium. We observed that during differentiation BIN1 expression is conveyed with an increase in phospho-ezrin expression (Fig. 5D-E). Furthermore, BIN1 appears colocalized with phospho-ezrin on filopodia both under proliferating and differentiation conditions (Fig. 5F). Thus, pointing out that the expression and membrane localization of BIN1 and the active form of ezrin must be finely regulated during myoblasts differentiation.

As a central protein linking the actin cytoskeleton to the inner leaflet of the plasma membrane, ezrin is an important regulator of the mechanical cohesion of the membrane/actin interface (Diz-Muñoz et al., 2010; Sens and Plastino, 2015; Tsai et al., 2018). We thus assessed the cortex-to-membrane mechanics from the force required to form membrane tethers on myoblasts, as previously described (Diz-Muñoz et al., 2010). To this end, we used atomic force microscopy (AFM) cantilevers coated with poly-L-lysine to pull membrane tethers from the plasma membrane of control and *Bin1* knock-down C2C12 cells (Fig. 5G). We measured the static tether force, *f_0_*, which is required to hold a membrane tether at a constant height. This force depends on the bending stiffness of the membrane (κ), the in-plane membrane tension (σ), and the energy density of the membrane-to-cortex attachments (*W*_0_) (Sheetz, 2001):

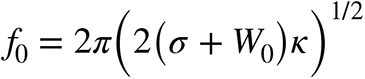

We found that the apparent static tether force is decreased in *Bin1* shRNA cells as compared to control conditions (Fig. 5G). In agreement, we also observed, using time-lapse phase-contrast videomicroscopy, the presence of prolonged retraction fibers similar to membrane tethers behind migrating *Bin1* knock-down cells (Fig. 5H). Therefore, suggesting that BIN1 regulates, either directly or through its association with ezrin, the stability of the plasma membrane/actin interface.

## Discussion

We unravel here an unexpected role of BIN1 in promoting filopodia formation in skeletal muscle cells, structures that play a crucial role in myoblast adhesion and fusion in mammalian cells and *Drosophila* (Segal et al., 2016; Zhang et al., 2020). The sequence of events that can be envisioned for the BIN1-mediated formation of filopodia at the myoblast cortex is presented in Fig. 6.

**Figure 6.**
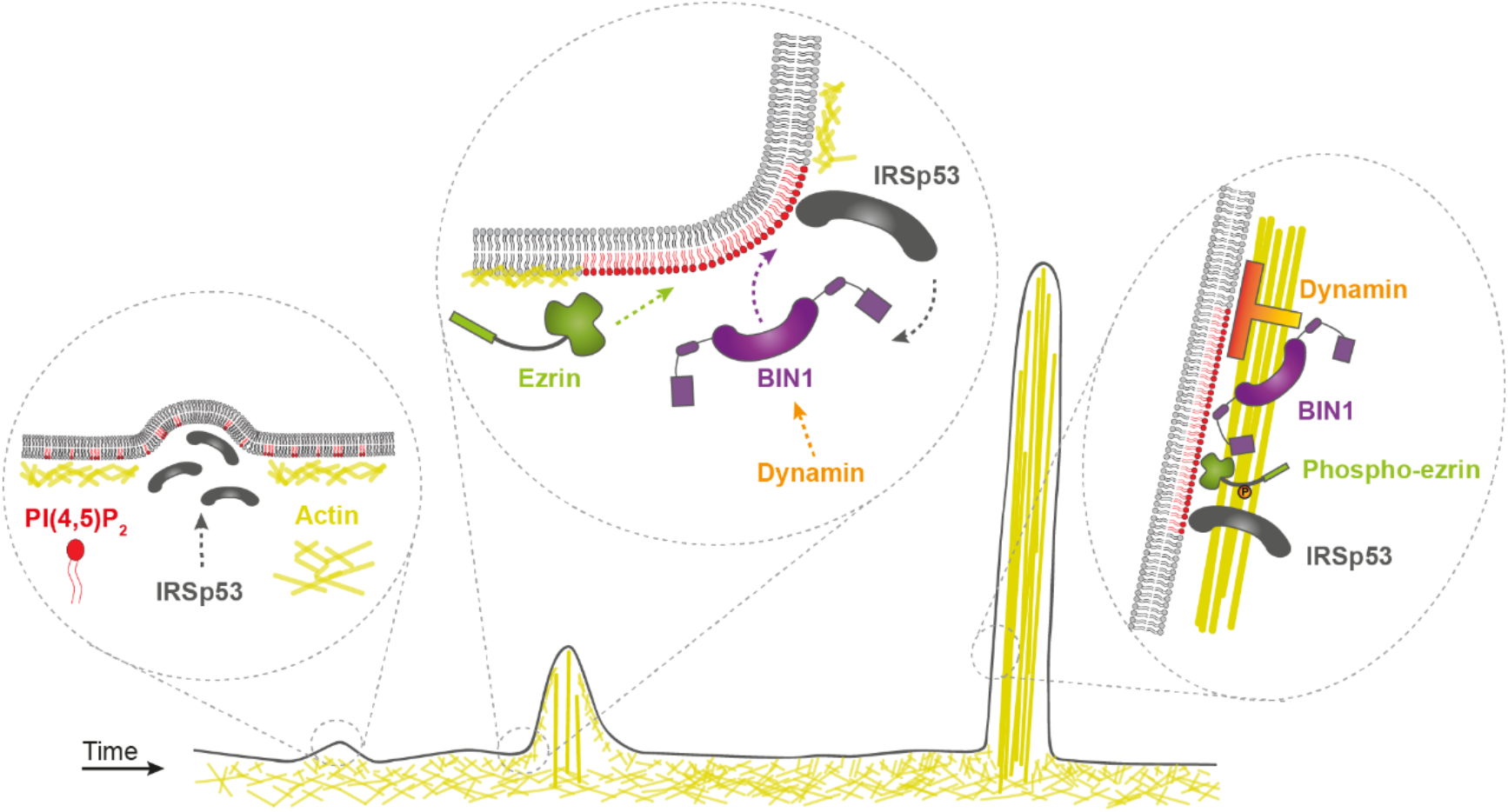
Model of BIN1-mediated filopodia formation at the myoblast membrane. The following steps are shown: 1) formation of BIN1-mediated filopodia-like structures requires an IRSp53-based actin complex leading to the initial evagination of the plasma membrane (Prévost et al., 2015). 2) Binding of BIN1 at the plasma membrane might led to the formation of a PI(4,5)P2-rich interface (Picas et al., 2014). This PI(4,5)P2 enrichment might, in turn, facilitate the accumulation of PI(4,5)P2-interacting proteins involved in the formation of actin-rich protrusions such as IRSp53, ezrin, and dynamin at the cell cortex. 3) The actin bundling ability of BIN1 and its association with phospho-ezrin, IRSp53 and dynamin (Zhang et al., 2020) facilitates its localization during filopodia formation. The different molecular players involved in this process are: IRSp53 in dark gray, BIN1 in magenta, dynamin in orange, ezrin and phospho-ezrin in green, actin in yellow, and PI(4,5)P2 in red.

Whereas BIN1 was shown to promote positive membrane curvature (Lee et al., 2002; Picas et al., 2014), we show here that BIN1 could also regulate the formation of membrane protrusions that have a negatively curved membrane geometry. Indeed, BIN1 expression in skeletal muscle cells promotes filopodia formation, and, reciprocally, its knock-down decreases their density at the cell surface, including at intercellular cell-cell contacts. High-resolution microscopy revealed that BIN1 is present at the base and all along the filopodia. Such an unexpected function was also reported in the case of PACSIN2/Syndapin II, an F-BAR domain protein shown to be involved in cellular protrusion formation (Qualmann and Kelly, 2000), presumably through the binding to the positively curved membrane at the base and the tip of filopodia (Shimada et al., 2010).

Our proteomic analysis revealed that BIN1 is associated with proteins having important functions during filopodia formation such as actin, dynamin, IRSp53, and the ERM proteins. We demonstrated that the generation of these filopodia by BIN1 requires IRSp53, an I-BAR protein that initiates the formation of filopodia (Lim et al., 2008; Disanza et al., 2013; Prévost et al., 2015) and is implicated in the generation and maintenance of filopodia during the fusion process in *Drosophila* (Segal et al., 2016). IRSp53 acts as a membrane-curvature sensing platform for assembling protein complexes (Bisi et al., 2020). Our *in vitro* data with purified IRSp53, which is active in this condition in the absence of Cdc42 (Tsai et al., 2022), show that IRSp53 allows BIN1 recruitment at the base of negatively-curved membranes. BAR domain proteins are well known to generate stable lipid membrane microdomains (Zhao et al., 2013; Picas et al., 2014). BIN1 has a PI domain that targets it to membrane domains, and we previously showed that BIN1 induces the clustering of PI(4,5)P_2_. IRSp53 directly binds PI(4,5)P_2_-rich membrane and deforms them (Mattila et al., 2007), and might also promote the formation of PI(4,5)P_2_ clusters (Zhao et al., 2013). We thus could envision a positive feedback loop with both proteins favoring their respective recruitment at specific membrane sites, where IRSp53 recruitment, in turn, assists BIN1 localization at the base of filopodial structures. Moreover, these PI(4,5)P_2_ clusters might facilitate the recruitment of proteins having PI(4,5)P_2_ binding motifs, such as actin regulators involved in protrusion formation (Senju et al., 2017; Senju and Lappalainen, 2019). The recruitment of actin regulators could also be directly mediated by BIN1 and IRSp53-dependent protein-protein interactions. Among the BIN1 partners known to regulate actin remodeling are WASP and dynamin (Yamada et al., 2009; Chuang et al., 2019; Zhang et al., 2020). Moreover, the SH3 domain of IRSp53 allows it to interact with VASP (Disanza et al., 2013), N-WASP (Lim et al., 2008), WAVE and mDia (Goh et al., 2012) or also EPS8, an actin-filament bundling and capping protein (Disanza et al., 2006; Hertzog et al., 2010).

Actin filament-bundling and membrane interactions are both essential features for the shape maintenance of filopodium (Mattila and Lappalainen, 2008; Svitkina, 2018). BIN1 could participate in F-actin bundling at least *via* two mechanisms. We discovered through *in vitro* assays a direct role of BIN1 in F-actin bundling, that combined with its direct association with actin, explains why BIN1 is closely connected with F-actin along filopodia. Moreover, the interaction of BIN1 with its downstream partner dynamin (Kojima et al., 2004; Royer et al., 2013) is likely to enhance the formation of actin-rich protrusions *via* its multifilament actin-bundling ability required for efficient myoblast fusion (Zhang et al., 2020). Furthermore, our proteomic analysis of the BIN1 partners confirmed its association with dynamin and actin and, importantly, identified the ERM family proteins. BIN1 is the only amphiphysin member tested to associate with ezrin, to our knowledge. We focused on the BIN1/ezrin interaction and function because ezrin is a membrane-actin linker that anchors F-actin to the plasma membrane at different types of membranes protrusions (Fehon et al., 2010), and appears enriched in filopodia (Osawa et al., 2009).We showed that BIN1 and the phosphorylated active version of ezrin are co-localized in filopodia, and in the absence of BIN1, phosphorylated ezrin is no more enriched in filopodia. BIN1 could participate in ezrin recruitment through several non-exclusive mechanisms. First, BIN1 SH3 domain is crucial for its association with ezrin, suggesting direct protein-protein interaction. Secondly, ezrin binding could also be mediated by the IRSp53 protein which was shown to enrich ezrin on negatively curved membranes (Tsai et al., 2018). Local depletion of ezrin from the plasma membrane is required to initiate actin-driven protrusion formation (Welf et al., 2020). Our study shows that BIN1-mediated filopodia formation is likely to occur once the protrusion is initiated, as it requires IRSp53 for its localization at negatively-curved membrane indentations. Thus, BIN1/ezrin association might enhance the membrane/actin interaction at later stages of filopodia formation. This is in agreement with the described role of ezrin as a major regulator linking the plasma membrane to the cortical actin while allowing the rapid remodeling of both structures (Diz-Muñoz et al., 2010; Rouven Brückner et al., 2015; Korkmazhan and Dunn, 2022).

The membrane-to-cortex architecture is tightly tuned in dynamic processes such as membrane protrusion formation during cell migration or membrane fusion in myo^-^ blasts (Diz-Muñoz et al., 2010; Kim et al., 2015; Sens and Plastino, 2015; Chakraborty et al., 2022). Using AFM, we identified that BIN1 regulates the apparent static tether force. The absence of BIN1 would contribute to a disorganization of the PI(4,5)P_2_ /ezrin interface at the plasma membrane and consequently, to a loose membrane-to-cortex attachment and a decrease in the formation of filopodia. Indeed, the long membrane tails in migrating cells that we observed upon BIN1 knock-down would reflect a loosening of membrane-cortex attachments. Furthermore, all the BIN1-mediated processes here identified (F-actin bundling) and partners (IRSp53, dynamin) are mechanisms previously identified to participate in the formation of membrane protrusion required for myoblast fusion in mammals and insects (Vasyutina et al., 2009; Bach et al., 2010; Sens et al., 2010; Shilagardi et al., 2013; Bothe et al., 2014; Segal et al., 2016; Zhang et al., 2020). Accordingly, we confirmed that BIN1 is required for myoblast fusion in C2C12 myoblasts, as previously reported *in cellulo* (Wechsler-Reya et al., 1998) and *in vivo* (Lee et al., 2002; Fernando et al., 2009; Prokic et al., 2020). BIN1 isoform 8 appears dispensable for muscle development but is required for muscle regeneration in adulthood (Prokic et al., 2020) and the D151N or the ΔSH3 BIN1 mutants are associated with the autosomal recessive form of centronuclear myopathies (Nicot et al., 2007; Royer et al., 2013).

In conclusion, we discovered a novel and unexpected role for BIN1 in the formation of filopodia, adding complexity in the diversity of the molecular mechanisms leading to filopodia formation (Dobramysl et al., 2021). Our study suggests a dual function of BIN1 both as a scaffold to recruit proteins, including ezrin, at the cell cortex and as an actin-bundling protein. We found a tandem role of IRSp53 and BIN1, in filopodia formation. The association of IRSp53 with antagonistic BAR domain proteins was reported to participate in filopodia formation (Galic et al., 2014; Dobramysl et al., 2021) or to assist endocytosis (Veltman et al., 2011; Bisi et al., 2020) in different cell types and organisms. Why proteins with opposite curvature domains are required to enable distinct remodeling events at the plasma membrane is yet an open question but suggests that the spatio-temporal regulation of these events is a complex molecular process.

## Methods

### Reagents

Natural and synthetic phospholipids, including POPS, POPC, POPE, brain total lipid extract, brain L-α-phosphatidylinositol-4,5-bisphosphate, and TopFluor-TMR-PI(4,5)P_2_ are from Avanti Polar Lipids. OG-DHPE and β-casein from bovine milk (>98% pure) were from Sigma-Aldrich. Alexa Fluor 647 and 488 Maleimide labelling kits are from Invitrogen. Culture-Inserts 2 Well for self-insertion were purchased from ibidi.

The following antibodies were used in this study: monoclonal mouse antibody to BIN1 (clone C99D against exon 17) from Millipore, polyclonal rabbit antibody to IRSp53 (BAIAP2) from Atlas Antibodies (HPA023310), HRP-conjugated beta actin monoclonal antibody from Proteintech (HRP-60008), ezrin antibody from M. Arpin laboratory(Algrain et al., 1993), monoclonal rabbit antibody to phospho-Ezrin (Thr567)/Radixin (Thr564)/Moesin (Thr558) Cell Signaling (Cat. 3726). Atto390 488 phalloidin was from Sigma.

### Constructs

pGEX-Sumo vector coding for GST-BIN1 isoform 8 was obtained by Gibson assembly from (Picas et al., 2014). pGEX vectors coding for GST-BIN1 isoform 8 ΔSH3 and D151N were obtained as in (Picas et al., 2014). 6xHis-Sumo ezrin and T567D vectors were obtained as in (Tsai et al., 2018). pEGFP-BIN1 D151N, pEGFP-BIN1 ΔSH3 were obtained as in (Nicot et al., 2007). pEGFP-BIN1 isoform 8 was obtained from P. De Camilli (Yale University, New Haven). pEYFP-BIN1 isoform 1 and pEGFP-Amphiphysin1 were obtained from L. Johannes (Institut Curie UMR144, Paris). pEGFP-dynamin2 was a gift from Sandra Schmid (Addgene plasmid # 34686). mCherry-ezrin WT and T567 was obtained from (Gautreau et al., 2000) after cloning into pENTR1A Gateway entry vector (Invitrogen) and recombined into mCherry-C1. full-length human IRSp53 (UniProt, no. Q9UQB8; Homo sapiens) was subcloned into the pGEX-6P-1 vector (Cytiva), as previously reported(Tsai et al., 2022). All the plasmids used in this study (Table 1) were sequenced.

**Table 1.**
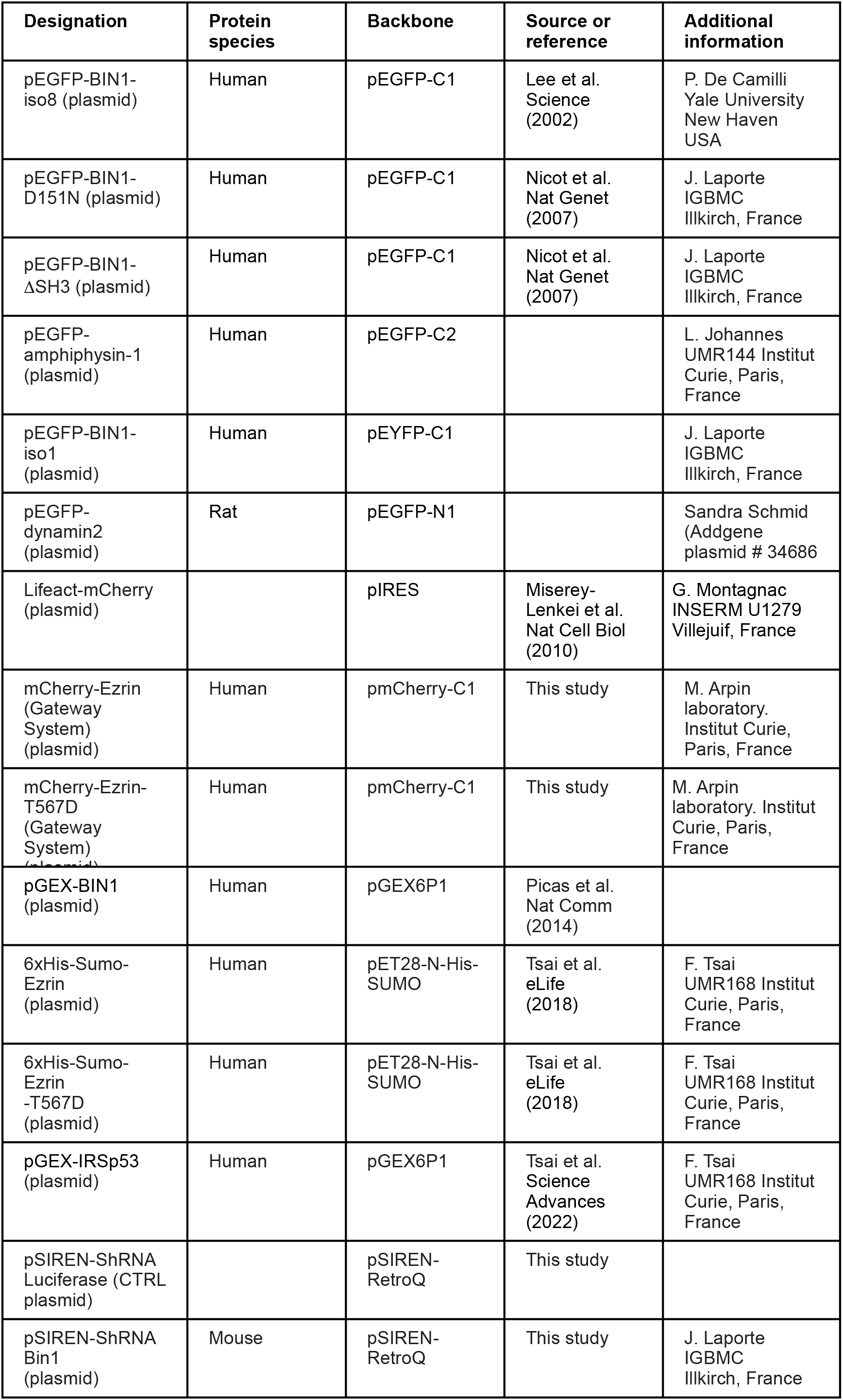
Summary of the plasmids used in this study.

| **Designation** | **Protein species** | **Backbone** | **Source or reference** | **Additional information** |
| --- | --- | --- | --- | --- |
| pEGFP-BIN1-iso8 (plasmid) | Human | pEGFP-C1 | Lee et al. Science (2002) | P. De Camilli<br>Yale University<br>New Haven<br>USA |
| pEGFP-BIN1-D151N (plasmid) | Human | pEGFP-C1 | Nicot et al. Nat Genet (2007) | J. Laporte<br>IGBMC<br>Illkirch, France |
| pEGFP-BIN1-ΔSH3 (plasmid) | Human | pEGFP-C1 | Nicot et al. Nat Genet (2007) | J. Laporte<br>IGBMC<br>Illkirch, France |
| pEGFP-amphiphysin-1 (plasmid) | Human | pEGFP-C2 |  | L. Johannes<br>UMR144 Institut<br>Curie, Paris,<br>France |
| pEGFP-BIN1-iso1 (plasmid) | Human | pEYFP-C1 |  | J. Laporte<br>IGBMC<br>Illkirch, France |
| pEGFP-dynamin2 (plasmid) | Rat | pEGFP-N1 |  | Sandra Schmid<br>(Addgene<br>plasmid # 34686) |
| Lifeact-mCherry (plasmid) |  | pIRES | Miserey-Lenkei et al. Nat Cell Biol (2010) | G. Montagnac<br>INSERM U1279<br>Villejuif, France |
| mCherry-Ezrin (Gateway System) (plasmid) | Human | pmCherry-C1 | This study | M. Arpin<br>laboratory.<br>Institut Curie,<br>Paris, France |
| mCherry-Ezrin-T567D (Gateway System) (plasmid) | Human | pmCherry-C1 | This study | M. Arpin<br>laboratory. Institut<br>Curie, Paris,<br>France |
| pGEX-BIN1 (plasmid) | Human | pGEX6P1 | Picas et al. Nat Comm (2014) |  |
| 6xHis-Sumo-Ezrin (plasmid) | Human | pET28-N-His-SUMO | Tsai et al. eLife (2018) | F. Tsai<br>UMR168 Institut<br>Curie, Paris,<br>France |
| 6xHis-Sumo-Ezrin-T567D (plasmid) | Human | pET28-N-His-SUMO | Tsai et al. eLife (2018) | F. Tsai<br>UMR168 Institut<br>Curie, Paris,<br>France |
| pGEX-IRSp53 (plasmid) | Human | pGEX6P1 | Tsai et al. Science Advances (2022) | F. Tsai<br>UMR168 Institut<br>Curie, Paris,<br>France |
| pSIREN-ShrRNA Luciferase (CTRL plasmid) |  | pSIREN-RetroQ | This study |  |
| pSIREN-ShrRNA Bin1 (plasmid) | Mouse | pSIREN-RetroQ | This study | J. Laporte<br>IGBMC<br>Illkirch, France |

### Cell culture

HeLa cells (CCL2 from ATCC) were cultured in DMEM medium (Gibco BRL) supplemented with 10% fetal bovine serum, 100 U/ml penicillin/streptomycin, and 2 mM glutamine. C2C12 mouse myoblasts (ATCC CRL-1772) were grown in DMEM/ Ham’s F-12 (1:1) supplemented with 10% fetal bovine serum. To induce differentiation, the growth medium was replaced with differentiation medium consisting of DMEM/Ham’s F-12 supplemented with 2% fetal calf serum (Hyclone/Perbio Sciences, Brebieres, France). All cells were tested mycoplasma free.

### Co-immunoprecipitation, western blot experiments and proteomic assays

To test the interaction between Amphiphysin isoforms and endogenous actin or ERM proteins, HeLa cells (CCL2 from ATCC) were seeded on 50 cm petri dishes overnight. Cells were then transfected for 24 h with either GFP, full-length GFP-BIN1 isoform 8, GFP-BIN1 isoform 1 and GFP-Amphiphysin1 using x-tremGENE9 (Roche). Cells were then trypsinized, washed once in PBS, and incubated on ice for 60 min in a lysis buffer: 25 mM Tris pH 7.5, 50–100 or 200 mM NaCl, and 0.1% NP40. Cells were then centrifuged 10 min at 10,000×g to collect the supernatant. Extracts were processed for co-immunoprecipitation using GFPTrap beads (Chromotek) (20µl GFPTrap beads per condition) for 3h at 4°C in lysis buffer. Beads were washed four times in lysis buffer and then processed for western-blotting.

To test the interaction of BIN1 and its mutated variants, C2C12 myoblasts were seeded overnight and transfected for 24h with either GFP, full-length GFP-BIN1 isoform 8 WT, ΔSH3 and D151N mutants using JetPEI (Ozyme). Co-immunoprecipitation assay was performed as detailed above.

For western-blot experiments cell lysis was performed in 25 mM Tris pH 7.5, 50 mM NaCl, 0.1% NP40, and a protease inhibitor cocktail (Sigma). The following primary antibodies were used: rabbit anti-ezrin (from M. Arpin laboratory (Algrain et al., 1993); 1:1000), mouse anti-actin (Sigma; 1:1000), mouse anti-BIN1 (clone C99D from Millipore; 1:1000), phosphor-ezrin. Secondary Horseradish Peroxidase (HRP)-coupled antibodies were from Jackson Laboratories.

For proteomic analysis, HeLa cells were transiently transfected with either GFP or GFP-BIN1 using X-tremeGENE 9 (Sigma-Aldrich), according to the manufacturer’s instructions. HeLa cells, with high transfection efficiency, were used because we previously developed and set up proteomic analysis using this cell line and to ensure that cell type specificities would not bias the obtained BIN1 interactors. Cell lysis and coimmunoprecipitation using GFP-Trap (Chromotek) was performed as described above. Mass spectrometry analysis was performed at the Proteomic platform of Institut Jacques Monod (Paris, France). Positive hits binding to GFP-Bin1 were selected relatively to their corresponding Mascot score.

### Short interfering RNA (shRNA)

shRNA constructs were engineered on a pSIREN retroviral vector (Clontech). To deplete the endogenous expression of BIN1, the oligonucleotide 5’-GATCCG**CCT-G A T A T C A A G T C G C G C A T T** T T C A A G A G A **A A T G C G C-GACTTGATATCAGG**CTTTTTTACGCGTG-3’ was inserted into pSIREN. Bold letters correspond to exons 5/6’s junction of mouse *Bin1*. As a control, we used the oligonucleotide 5’-GTTGCGCCCGCGAATGATATATAATGttcaagagaCATTATATATCATTCGCGGGCGCAAC-3’ sequence against luciferase.

HEK293T cells expressing shRNA *Bin1* or shRNA luciferase (CTRL) were cultured, and cell-free supernatants containing retrovirus were harvested. Two sequential transductions of C2C12 myoblasts were performed, as previously described (Meriane et al., 2000), by adding 2 mL of filtered supernatant (0.45 mm PES sterile syringe filter) within a time window of 6h between transductions. Double transduced cells were kept for 72h in culture. After this time, cells were kept under puromycin at 1 μg/ml for 72h before performing the experiments. For each cycle of double transient transduction, shRNA efficiency was determined by western blotting showing to ensure a complete depletion of the BIN1 protein. Under these experimental conditions, we did not observe significant cellular mortality due to puromycin selection, suggesting that the entire population of cells should be infected by the shRNA.

To ensure that defects in cell proliferation and motility in the shRNA *Bin1* cell line, shRNA BIN1 C2C12 were seeded at higher density (i.e., at the onset of differentiation).

### RNA interference (siRNA)

The siRNA used in this study to IRSp53 was ON-TARGET plus siIRSp53 (BAIAP2) mouse (Horizon Discovery, Cat# J-046696-11) (Rodríguez-Pérez et al., 2021). The siRNA sequence targeting luciferase (CGUACGCGGAAUACUUCGA) was used as a control and was obtained from Sigma. siRNA delivery was performed using Lipofectamine RNAiMAX, according to the manufacturer’s instructions.

### Time-lapse fluorescence microscopy

C2C12 cells were seeded on FluoroDish (WPI, France) cell culture dishes overnight. Cells were then transfected for 12 h with either GFP-BIN1, D151N or ΔSH3 mutants and Lifeact-mCherry; GFP or GFP-BIN1 and its mutated variants and mCherry-ezrin using JetPEI (Ozyme) following the manufacturer’s instructions. Live-cell imaging was performed on a Spinning disk microscope based on a CSU-X1 Yokogawa head mounted on an inverted Ti-E Nikon microscope equipped with a motorized XY Stage. Images were acquired through a 60x objective NA 1.4 Plan-Apo objective with a Photometrics Coolsnap HQ2 CCD camera. Optical sectioning was performed using a piezo stage (Mad City Lab). A dual Roper/ Errol laser lounge equipped with 491 and 561 nm laser diodes (50 mW each) and coupled to the spinning disk head through a single fiber was used. Multi-dimensional acquisitions were performed in streaming mode using Metamorph 7 software. Images were collected every second (500 msec exposure) during 60s.

### Immunofluorescence microscopy

Fixed cells were obtained after incubation with 3.2% paraformaldehyde (PFA) for 10 min at room temperature, washed with PBS, incubated with PBS-0.1 M NH_4_Cl for 5 min and then washed with PBS. Finally, cells were permeabilized in 0.1% Triton X-100 for 3 min and blocked with 1% BSA during 10 min. Fixed cells were mounted using mowiol mounting agent and visualized using a Leica DMRA and a CoolSnap-HQ2 camera, 100x objective NA 1.25 oil Ph 3 CS (HCX PL APO) and analyzed with the Metamorph software. 3D stacks were acquired and deconvolved to build a projection on one plane using Image J.

For myotube imaging, images were acquired on a Zeiss LSM880 Airyscan confocal microscope (MRI facility, Montpellier). Excitations sources used were: 405 nm diode laser, an Argon laser for 488 nm and 514 nm and a Helium/Neon laser for 633 nm. Acquisitions were performed on a 63x/1.4 objective. Multidimensional acquisitions were performed via an Airyscan detector (32-channel GaAsP photomultiplier tube (PMT) array detector).

Images are presented as a z-projection of all planes.

### Tether pulling experiments

Tether pulling experiments were performed on a JPK Nanowizard III mounted on an inverted Zeiss wide-field microscope. Olympus Biolevers (k = 6 mN·m^-1^) were cleaned in acetone for 5 minutes and then plasma-cleaned for 10 min. Then, cantilevers were soaked briefly in 0.1 M of NaHCO_3_ (pH 9.0), air dried and immersed in 0.01% poly-L-lysin overnight at 4°C in a humid chamber. Before the measurements, cantilevers were rinsed three times in PBS and mounted on the AFM cantilever holder. The cantilever spring constant was determined by the thermal noise method, as detailed in (Schillers et al., 2017). For the measurement, cells seeded for 24h on FluoroDish cell culture dishes were kept on DMEM-F12 medium at 37°C and not used longer than 1 h for data acquisition. Static tether force measurements were performed by retracting the cantilever for 6 μm at a speed of 10 μm·s^-1^, and the position was kept constant for 30 s. Resulting force–time curves were analyzed using the JPK analysis software. Tether force values represent the average value obtained from tether pulling maps performed over the whole apical membrane of adherent C2C12 cells after applying 3 x 3 grids of 10 μm^2^.

### Protein purification and fluorescent labelling

Recombinant human full-length BIN1 isoform 8, Amphiphysin 1, ezrin wild-type and ezrin T567D were expressed in Rosetta 2 bacteria and purified by affinity chromatography using glutathione Sepharose 4B beads as previously published (Picas et al., 2014; Tsai et al., 2018). Recombinant proteins were labelled by conjugation with either Alexa Fluor 488 or 647 following maleimide chemistry (Invitrogen), as in (Picas et al., 2014; Tsai et al., 2018).

Recombinant human full-length IRSp53 (Uniprot #Q9UQB8) was purified and labeled with Alexa Fluor 488 as previously described(Tsai et al., 2022).

Muscle actin was purified from rabbit muscle and isolated in monomeric form in G-buffer (5 mM Tris-Cl-, pH 7.8, 0.1 mM CaCl2, 0.2 mM ATP, 1 mM DTT, 0.01% NaN3) as previously described (Spudich and Watt, 1971).

### Lipid bilayer experiments

Supported lipid bilayers were prepared as described in (Braunger et al., 2013). The lipid mixture consisted of: 60% POPC, 20% POPE, 10-15% POPS and 5% PI(4,5)P_2_. The amount of total negatively charged lipids was kept to 20% for any of the mixtures containing PIs at the expenses of brain-phosphatidylserine. Fluorescent TopFluor-TMR-PI(4,5)P_2_ was added to 0.1%. Experiments were performed by injecting 15 µL of buffer (10 mM Tris, pH 7.4, 100 mM NaCl and 0.5 mg·ml^-1^ of casein). Supported lipid bilayers were imaged on a Zeiss LSM880 Airyscan confocal microscope (MRI facility, Montpellier). Excitations sources used were: Argon laser for 488 nm and 514 nm and a Helium/Neon laser for 633 nm. Acquisitions were performed on a 63x/1.4 objective.

### GUV preparation and observation

The lipid mixture used contain total brain extract supplemented with 5 mol% PI(4,5)P_2_ at 0.5 mg/mL in chloroform. If needed, the lipid mixture is further supplemented with 0.5 mol% BODIPY TR ceramide or 0.5 mol% OG-DHPE.

GUVs were prepared by the polyvinyl alcohol (PVA) gel-assisted formation method as previously described(Weinberger et al., 2013). Briefly, a PVA gel solution (5 %, w/ w, dissolved in 280 mM sucrose and 20 mM Tris, pH 7.5) warmed up to 50 °C was spread on clean coverslips (20mm × 20mm). The coverslips were cleaned by ethanol and then ddH_2_O twice. The PVA-coated coverslips were incubated at 50 °C for 30 min. Then, around 5 μl of the lipid mixture was spread on the PVA-coated coverslips, followed by placing them under vacuum at room temperature for 30 min. The covers^-^ lips were then placed in a petri dish and around 500 μl of the inner buffer was pipetted on the top of the coverslips. The inner buffer contains 50 mM NaCl, 20 mM sucrose and 20 mM Tris-HCl pH 7.5. The coverslips were kept at room temperature for at least 45 min, allowing GUVs to grow. Once done, we gently “ticked” the bottom of the petri dish to detach GUVs from the PVA gel. The GUVs were collected using a 1 ml pipette tip with its tip cut to prevent breaking the GUVs.

For all experiments, coverslips were passivated with a β-casein solution at a concentration of 5 g.L^-1^ for at least 5 min at room temperature. Experimental chambers were assembled by placing a silicon open chamber on a coverslip.

GUVs were first incubated with IRSp53 in the outer buffer (60 mM NaCl and 20 mM Tris-HCl pH 7.5) for at least 15 min at room temperature before adding BIN1 into the GUV-IRSp53 mixture. The final GUV-protein mixture was then incubated at least 15 min at room temperature before observation. Samples were observed using a Nikon Eclipse Ti microscope equipped with Yokogawa CSU-X1 spinning disk confocal head, 100X CFI Plan Apo VC objective (Nikon) and a sCMOS camera Prime 95B (Photometrics).

### F-actin bundling assay

Actin (1 µM, non-labeled) was polymerized 1 hour in a buffer containing 100 mM KCl, 1 mM MgCl_2_, 0.2 mM EGTA, 0.2 mM ATP, 10 mM DTT, 1 mM DABCO, 5 mM Tris pH 7.5 and 0.01% NaN3 in the presence of BIN1 at different concentrations. Then, 5 µl of the protein mixtures was diluted 20 times (i.e., 50 nM of filamentous actin in the presence of BIN1) in the same buffer supplemented with 0.3% methylcellulose and 660 nM of Alexa Fluor 546-phalloidin. Samples were observed using TIRF microscopy (Eclipse Ti inverted microscope, 100x TIRF objectives, Quantem 512SC camera).

### Image processing and analysis

Protein binding was quantified by measuring the mean grey value of still confocal images of either AlexaFluor 488 ezrin, WT or T567D, in the absence of BIN1 or Amphiphysin 1. The obtained average intensity was then used to estimate the fold increase in the binding of ezrin, WT or T567D, in the presence of 0.1 μM of BIN1 or Amphiphysin 1 tagged with AlexaFluor 647. Each data set was performed using the same supported lipid bilayer preparation and confocal parameters were kept constant between experiments and samples. Mean gray values were measured once the steadystate of protein binding was reached, which was estimated to be ≥ 600 s. Mean gray values were measured using Image J (Schindelin et al., 2012).

To obtain actin fluorescence intensities on a filament or bundles, we manually defined a ROI, a 6 pixel-width line perpendicularly to the filament or bundle. We then obtained the intensity profile of the line in which the x-axis of the profile is the length of the line and the y-axis is the averaged pixel intensity along the width of the line. The actin intensity was the maximum intensity value in the intensity profile.

Filopodia density, morphology and dynamics were analyzed by a custom written program allowing the semi-automatic tracking the Life-actin signal of time-lapse movies using Image J.

### Statistical analyses

Results are shown as a mean ± standard deviation (s.d.). Unless stated otherwise, average values represent at least three technical replicates in the case of *in vitro* experiments and biological replicates in the case of cellular studies. Statistical significance was assessed by Welch’s t test, unless stated otherwise.

## Acknowledgements

The authors thank J. Viaud for kindly providing the PH-PLCd-Alexa647 probe. We thank Y. Senju (Okayama University, Japan) and Pekka Lappalainen (University of Helsinki, Finland) for IRSp53 purification and C. Le Clainche (Institute for Integrative Biology of the Cell, Gif-sur-Yvette, France) for actin purification. We also thank B. Cowling for the design of *Bin1* shRNA constructs. We also thank C. Cazevieille for assistance in TEM and J. Mateos-Langerak in SIM OMX imaging. The authors thank P. Sens for scientific discussions. The authors acknowledge the Nikon Imaging Center at Institut Curie-CNRS, the imaging facility MRI, the PICT-IBiSA, members of the national infrastructure France-BioImaging infrastructure supported by the French National Research Agency (ANR10-INBS-04). L.P. acknowledges the ATIP-Avenir program (AO-2016) for financial support. This project was also supported by grants from the Agence Nationale de la Recherche (ANR) (ANR-13-BSV2-0004-01), the Labex Cell(n)Scale (N° ANR-10-LBX-0038) part of the IDEX PSL (N° ANR-10-IDEX-0001-02 PSL), the Association Française contre les Myopathies (15352), the CNRS, the INSERM, the University of Montpellier, University of Strasbourg, the College de France, and the Institut Curie.

## Author contributions

Conceptualization of the study: L.P., B.G., C. G-R. and S.M-L. Performed experiments and data analysis: L.P., F.C., C.A-A, H.B., F-C-T., J. P., F.R., P.M., S.B., A-S. N, and S.M-L. Supervision: L.P., J.L., P.B., B.G., C.G-R., and S.M. Writing: L.P., C.G-R. and S.M-L with inputs from all authors.

## SUPPLEMENTARY INFORMATION

**Figure S1.**
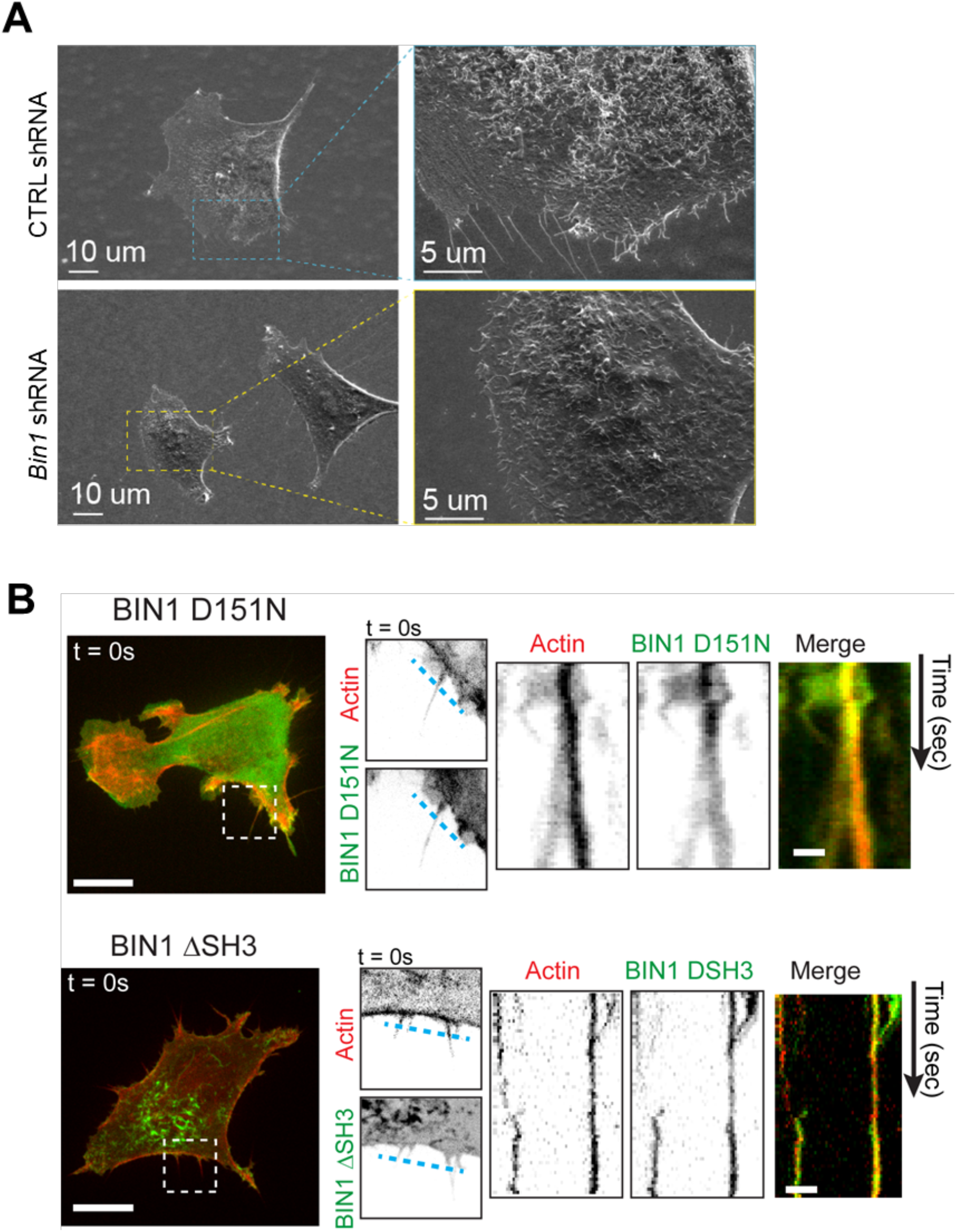
**A)** Representative SEM images displaying the plasma membrane organization of isolated C2C12 myoblasts expressing either *Bin1* shRNA or CTRL shRNA (*luciferase*) under proliferative conditions. **B)** Snap-shots of spinning disk images showing C2C12 cells co-transfected with either GFP-BIN1 D151N or GFP-BIN1 ΔSH3 mutant and Lifeact-mCherry (actin). Scale bar, 10 µm.

**Figure S2.**
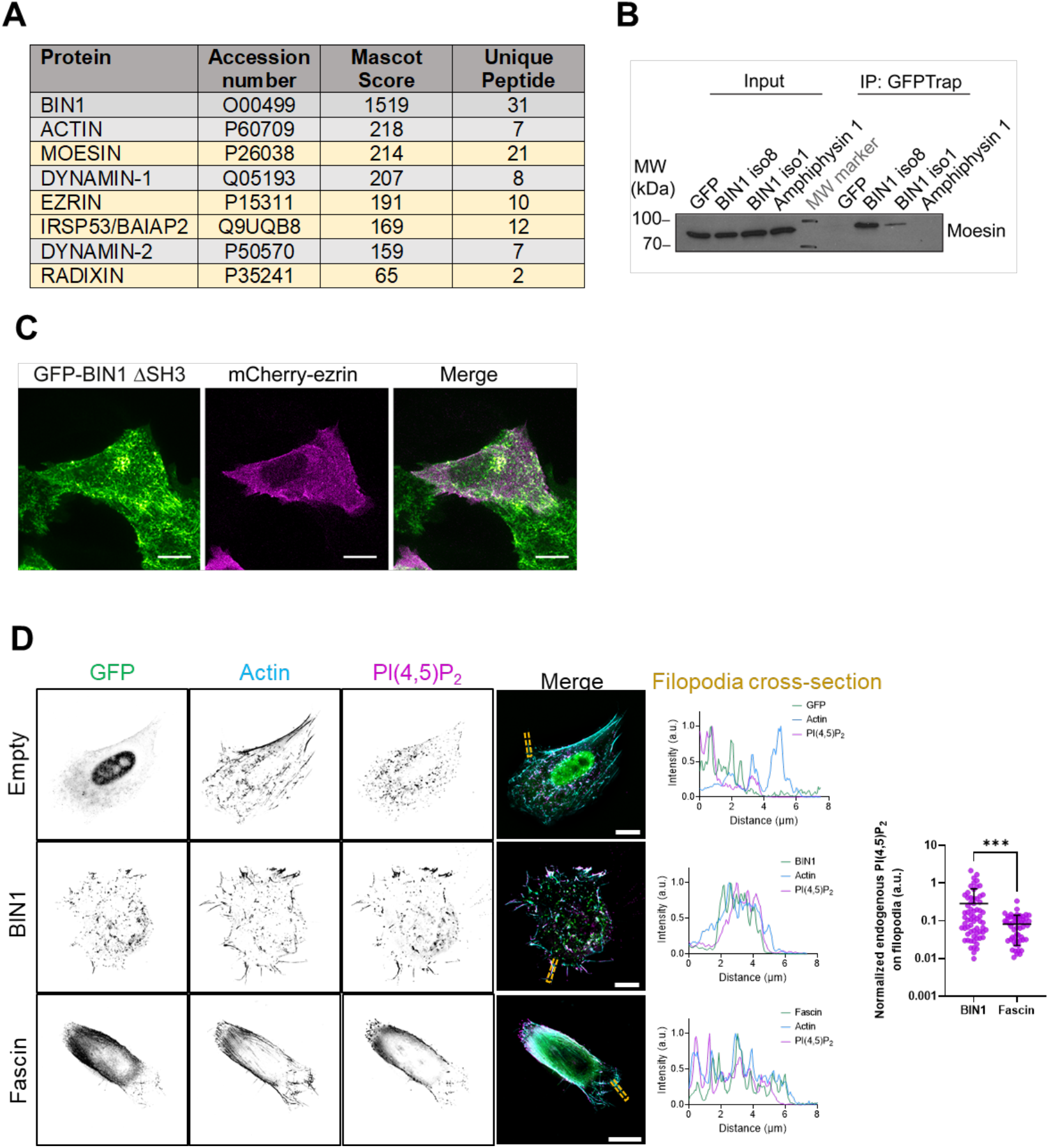
**A,** Identification of some binding partners of GFP-BIN1 using proteomic analysis. GFP alone was used as a negative control. Only Mascott Scores superior to 50 were considered as positive hits. In gray, already known BIN1 partners. In yellow, new identified BIN1 partners. **B)** GFP-Trap pull-downs from extracts of HeLa cells transfected with plasmids encoding GFP, GFP-tagged amphiphysin1, BIN1 iso1 and BIN1 iso8. Moesin was revealed by western-blotting. **C)** Time projection images from spinning disk movies (500 ms, 120 s) of C2C12 myoblasts co-expressing mCherry-ezrin (magenta) and GFP-BIN1 ΔSH3 (green). Scale bar, 10 µm. **D)** Wild-field deconvoluted images of C2C12 cells transfected with either GFP, GFP-BIN1 iso8 or GFP-Fascin and stained for endogenous PI(4,5)P2 (PH-PLCd-Alexa647, magenta) and F-actin (phalloidin, cyan). Cross-section analysis of filopodia structures corresponding to the highlighted yellow region. Quantification of the normalized intensity of endogenous PI(4,5)P2 (as the ratio between PI(4,5)P2 intensity at filopodia respect to PI(4,5)P2 intensity of the whole cell) at filopodia-like structures induced by BIN1 and Fascin, respectively. Scale bar, 10 µm.

**Figure S3.**
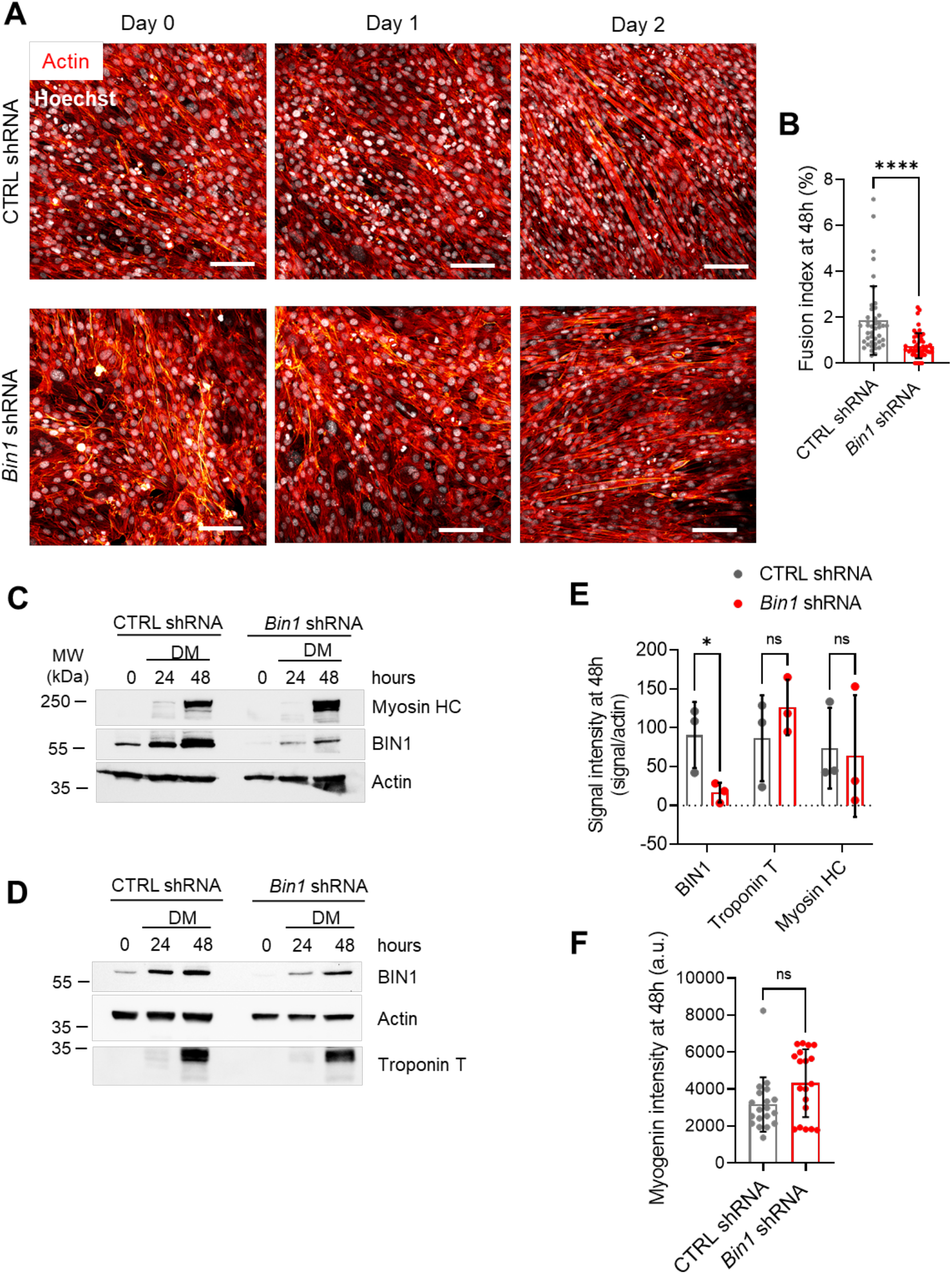
**A)** Confocal images of C2C12 myoblasts CTRL shRNA or *Bin1* shRNA stained for F-actin (phalloidin, in red) and the nuclei (Hoechst, in gray). Scale bar = 100 µm. **B)** Quantification of the fusion index (number of nuclei in myotubes divided by the total number of nuclei) of the C2C12 CTRL shRNA (gray) and *Bin1* shRNA (red) at 48h in differentiation medium. Number of myotubes, n = 43 and 50, for CTRL shRNA and *Bin1* shRNA, respectively. **C, D)** Western-blot analysis of the endogenous expression of actin, BIN1, myosin heavy-chain (HC), and Troponin T in CTRL shRNA and *Bin1* shRNA C2C12 myoblasts in growth medium (0 hours, undifferentiated) and grown in differentiation medium (DM) at 24h and 48h. **E)** Quantification of the signal intensity (signal/actin) of the endogenous expression of BIN1, Troponin T and myosin HC in CTRL shRNA and *Bin1* shRNA C2C12 myoblasts grown in DM for 48h obtained from western-blot analysis. **F)** Quantification of the intensity of the endogenous expression of myogenin in CTRL shRNA and *Bin1* shRNA C2C12 myoblasts grown in DM for 48h from confocal images (immunofluorescence). Number of images, n = 19 and 19, for CTRL shRNA and *Bin1* shRNA, respectively Mann Whitney test: n.s > 0.1, * P < 0.1., **** P < 0.0001. All data represents three independent experiments (N = 3).

